# Stochastic Neural Networks for Automatic Cell Tracking in Microscopy Image Sequences of Bacterial Colonies

**DOI:** 10.1101/2021.04.27.441677

**Authors:** Sorena Sarmadi, James J. Winkle, Razan N. Alnahhas, Matthew R. Bennett, Krešimir Josić, Andreas Mang, Robert Azencott

## Abstract

We describe an automated analysis method to quantify the detailed growth dynamics of a population of bacilliform bacteria. We propose an innovative approach to frame-sequence tracking of deformable-cell motion by the automated minimization of a new, specific cost functional. This minimization is implemented by dedicated Boltzmann machines (stochastic recurrent neural networks). Automated detection of cell divisions is handled similarly by successive minimizations of two cost functions, alternating the identification of children pairs and parent identification. We validate this automatic cell tracking algorithm using recordings of simulated cell colonies that closely mimic the growth dynamics of *E. coli* in microfluidic traps. On a batch of 1100 image frames, cell registration accuracies per frame ranged from 94.5% to 100%, with a high average. Our initial tests using experimental image sequences of *E. coli* colonies also yield convincing results, with a registration accuracy ranging from 90% to 100%.

## 1. Introduction

Technology advances have led to increasing magnitudes of data generation with increasing levels of precision [25, 32, 82, 90]. Data generation presently far outpaces data analysis, however, and drives the requirement for analyzing such large-scale data sets with automated tools [16, 35, 41, 48, 66, 87, 89]. The main goal of the present work is to develop computational methods for an automated analysis of microscopy image sequences of colonies of *E. coli* growing in a single layer. Such recordings can be obtained from colonies growing in microfluidic devices, and they provide a detailed view of individual cell-growth dynamics as well as population-level, inter-cellular mechanical and chemical interactions [5,6, 19, 28, 29, 65].

However, to understand both variability and lineage-based correlations in cellular response to environmental factors and signals from other cells requires the tracking of large numbers of individual cells across many generations. This can be challenging, as large cell numbers tightly packed in microfluidic devices can compromise spatial resolution, and toxicity effects can place limits on the temporal resolution of the recordings [33, 40]. One approach to better understand and control the behavior of these bacterial colonies is to develop computational methods that capture the dynamics of gene networks within single cells [5, 19, 46, 94]. For these methods to have a practical impact, one ultimately has to fit the models to the data, which allows us to infer hidden parameters (i.e., characteristics of the behavior of cells that cannot be measured directly). Image analysis and pattern recognition techniques for biological imaging data [26, 43, 66], like the methods developed in the present work, can be used to track lineages and thus automatically infer how gene expression varies over time. These methods can serve as an indispensable tool to extract information to fit and validate both coarse and detailed models of bacterial population, thus allowing us to infer model parameters from recordings.

Here we describe an algorithm that provides *quantitative* information about the population dynamics, including the life cycle and lineage of cells within a population from recordings of cells growing in a mono-layer. A typical sequence of frames of cells growing in a microfluidic trap is shown in Fig. 1. We describe the design and validation of algorithms for tracking individual cells in sequences of such images [5, 46, 55]. After segmentation of individual image frames to identify each cell, tracking individual cells from frame to frame is a combinatorial problem. To solve this problem we take into account the unknown cell growth, cell motion, and cell divisions that occur between frames. Segmentation and tracking are complicated by imaging noise and artifacts, overlap of bacteria, similarity of important cell characteristics across the population (shape; length; and diameter), tight packing of bacteria, and large interframe durations resulting in significant cell motion, and up to a 30% increase in individual cell volume.

**Fig. 1.**
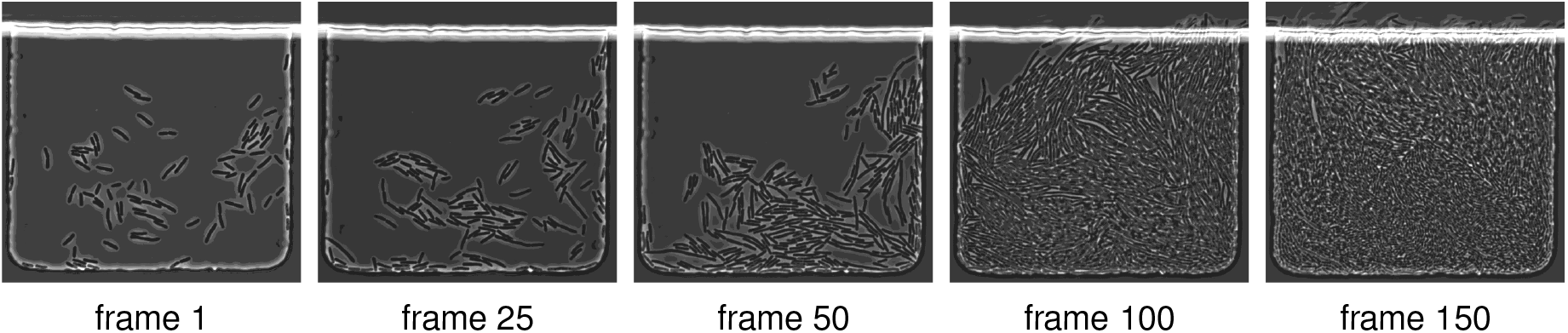
Typical microscopy image sequence. We show five frames out of a total of 150 frames of an image sequence showing the dynamics of E. coli in a microfluidic device [5].

### 1.1. Related Work

The present work focusses on tracking *E. coli* in time series of images. A comparison of different cell-tracking algorithms can be found in [35, 89]. Tracking and object recognition in time series of images is a challenging task that arises in numerous applications in computer vision [63, 97]. In image processing, motion tracking is often referred to as “*image registration*” [35, 57, 51, 60, 62, 61] or “*optical flow*” [23, 39, 47, 50, 95].

Our tests have shown that off-the-shelf image intensity driven techniques fail to provide a robust bacteria cell registration in the tightly packed colonies of rod-shaped *E. coli* bacteria considered here. Moreover, we are not interested in tracking individual pixels but rather cells (i.e., rod-shaped, deformable shapes), while recognizing events of cell division and recording cell lineage.

One approach proposed in prior work to simplify the tracking task is to make the experimental setup more rigid by confining individual cell lineages to small tubes; the associated microfluidic device is called a “*mother machine*” [18, 42, 58, 71, 79, 85]. The microfluidic devices we consider here yield more complicated data as cells are allowed to move and multiply freely in two dimensions (constrained to a mono-layer). We refer to Fig. 1 for a typical sequence of experimental images considered in the present work.

Turning to methods that work on more complex biological cell imaging data, we can distinguish different classes of tracking methods. “*Model-based evolution methods*” operate on the image intensities directly. They rely on particle filters [8, 70, 84] or active contour models [7, 37, 45, 52, 93, 96]. These methods work well if the cells are not tightly packed. However, they may lead to erroneous results if the cells are close together, the inter-cellular boundaries are blurry, or the cells move significantly. Our work belongs to another class—the so called “*detection-and-association methods*” [17, 22, 20, 44, 74, 78, 88, 92, 98], which first detect cells in each frame and then solve the tracking problem/association task across successive frames. Doing so necessitates the segmentation of cells within individual frames. We refer to [91] for an overview of cell segmentation approaches. Deep learning strategies have been widely used for this task [4, 34, 59, 67, 72,78, 77, 86, 87, 98]. We consider a framework based on convolutional neural networks (**CNN**s). Others have also used CNNs for cell segmentation [3, 59, 69, 76, 77]. We omit a discussion of our segmentation approach, as we do not view it as our main contribution (see Sec. 1.2). To solve the tracking problem after the cell detection, many of the methods cited above use hand-crafted association scores based on the proximity of the cells and shape similarity measures [44, 22, 88, 98]. We follow this approach here. We note that we not only consider local association scores between cells but also include measures for the integrity of a cell’s neighborhood (i.e., “*context information*”).

Our method is tailored for tracking cells in tightly packed colonies of rod-shaped *E. coli* bacteria. This problem has been considered previously [17, 74, 87, 92]. However, we are not aware of any large-scale datasets that provide ground truth tracking data for these types of recordings, but note that there are community efforts for providing a framework for testing cell tracking algorithms [64, 89].* Works that consider these data are for example [8, 59, 69, 73, 72, 98]. The cells in this dataset have significantly different characteristics compared to those considered in the present work. As we describe below, our approach is based on distinct characteristics of the bacteria cells and, consequently, does not directly apply to these data. Therefore, we have developed our own validation and calibration framework (see Sec. 2).

[27, 54, 53, 56] considered graph-based matching strategies for global association. Similar to the methods described above, they used association scores for tracking. Individual cells are represented as nodes, and neighbors are connected through edges. This is similar to our approach in that we construct local neighborhood relations based on a (modified) Delauny triangulation. By using a graph-like structure, cell divisions can be identified by detecting changes in the topology of the graph [54, 53]. We tested a similar strategy, but came to the conclusion that we cannot reliably construct neighborhood networks between frames for which topology changes only occur due to cell division; the main issue we observed is that the significant motion of cells between frames can introduce additional topology changes in our neighborhood structure. Consequently, we decided to relax these assumptions.

Some recent works jointly solve the tracking and segmentation problem [8, 36, 73, 74, 72, 98]. Contrary to observations we have made in our data, these approaches rely (with the exception of [74]) on the fact that the tracking problem is inherent to the segmentation problem (“*tracking-by-detection methods*” [98]; see also [87]). That is, the key assumption made by many of these algorithms is that cells belonging to the same lineage overlap across frames (see also [20]). In this case, cell-overlap can serve as a good proxy for cell-tracking [98]. We note that in our data we cannot guarantee that the frame rate is sufficiently high for this assumption to hold.

[73, 69, 36] exploited machine learning techniques for segmentation *and* motion tracking. One key challenge here is to provide adequate training data for these methods to be successful. We here describe simulation-based techniques that can be extended to produce training data, which we use for parameter tuning [94].

The works that are most similar to ours are [17, 74, 92]. Similar to our approach, they perform a local search to identify the best cell-tracking candidates across frames. One key difference across these works are the matching criteria. Moreover, [17, 74] employ a local greedy-search, whereas we consider stochastic neural network dynamics for optimization. [92] construct score matrices within a score based neighborhood tracking method; an integer programming method is used to generate frame-to-frame correspondences between cells and the lineage map. Other approaches that consider linear programming to maximize an association score function for cell tracking can be found in [21, 20, 98]

### 1.2. Contributions

For image segmentation, we first apply two well-known, powerful variational segmentation algorithms to generate a large training set of correctly delineated single cells. We can then train a CNN dedicated to segmenting out each single cell. Using a CNN significantly reduces the runtime of our computational framework for cell identification. The frame-to-frame tracking of individual cells in tightly packed colonies is a significantly more challenging task, and is hence the main topic discussed in the present work. We develop a set of innovative automatic cell tracking algorithms based on the successive minimization of three dedicated cost functionals. For each pair of successive image frames, minimizing these cost functionals over all potential cell registration mappings poses significant computational and mathematical challenges. Standard gradient descent algorithms are inefficient for these discrete and highly combinatorial minimization problems. Instead, we implement the stochastic neural network dynamics of Boltzmann machines (**BM**), with architectures and energy functions tailored to effectively solve our combinatorial tracking problem. Our major contributions are: **i)** The design of a multi-stage cell tracking algorithm, that starts with a parent-children pairing step, followed by removal of identified parent-children triplets, and concludes with a cell to cell registration step. **ii)** The design of dedicated BM architectures, with several energy functions, respectively, minimized by true parent-children pairing and by true cell-to-cell registration. Energy minimizations are then implemented by simulation of BM stochastic dynamics. **iii)** The development of automatic algorithms for the estimation of unknown weight parameters of our BM energy functions, using convex-concave programming tools [2, 30, 81, 80]. **iv)** The evaluation of our methodology on synthetic and real image sequences of cell colonies. The massive effort involved in human expert annotation of cell colony recordings limits the availability of “*ground truth tracking*” data for dense bacterial colonies. We therefore first validated the accuracy of our cell tracking algorithms on recordings of simulated cell colonies, generated by the dedicated cell colony simulation software [94]. This provided us with ground truth frame-by-frame registration for cell lineages, enabling us to validate our methodology.

### 1.3. Outline

In Sec. 2 we describe the synthetic image sequence of cell colonies considered here as benchmarks for our cell tracking algorithms. In Sec. 3 we describe key cell characteristics involved in our cost functionals. We define valid cell registration mappings between successive image frames in Sec. 4. We outline how to automatically calibrate the weights of our various penalty terms in Sec. 5. Our algorithms for pairing parent cells with their children and for cell-to-cell registration are developed in sections Sec. 6 through Sec. 12. We present our main validation results on long image sequences (time series of images) in Sec. 13 and conclude with Sec. 14.

## 2. Benchmark: Synthetic Videos of Simulated Cell Colonies

To validate our cell tracking algorithms, we consider simulated image sequences of dense cell populations. We refer to [94] for a detailed description of this mathematical model and its implementation. The simulated cell colony dynamics are driven by an agent based model [94], which emulates live colonies of growing, moving, and dividing rod-like *E. coli* cells in a 2D microfluidic trap environment. Between two successive frames *J, J*_+_, cells are allowed to move until they nearly bump into each other, and to grow at multiplicative rate denoted *g*.*rate* with an average value of 1.05 per minute (plus/minus a small random perturbation). For a cell of length *L*_0_ at birth, cell division occurs at length *L*_div_ = 2*L*_0_ + *ε*, where *ε* is a small uniformly distributed random variable. When a bacterial cell *b* of length *L*_div_ divides into two cells *b*_1_ and *b*_2_, their lengths *L*_1_, *L*_2_ satisfy *L*_1_ + *L*_2_ = *L*_div_ and *L*_1_*/L*_div_ is a random number in [0.45, 0.55]. All these random variables are independent of each other.

The simulation keeps track of cell lineage, cell size, and cell location (among other parameters). The main output of each such simulation considered here is a binary image sequence of the cell colony with a fixed interframe duration. Each such synthetic image sequence is used as the sole input to our cell tracking algorithm. The remaining meta-data generated by the simulations are only used as ground truth to evaluate the performance of our tracking algorithms.

We consider two benchmark datasets, BENCH1 and BENCH6, of several *synthetic* image sequences of simulated cell colonies with *cell growth factor g*.*rate* = 1.05 per minute. The generated binary images are of size 600 × 600 pixels, with interframe durations of 1 minute for BENCH1, and of 6 minutes for BENCH6. The associated image sequences involve 100 to 500 frames each. In Fig. 2 we display an example of two simulated consecutive frames separated by 1 minute. To simplify our presentation and validation tests, we control our simulations to make sure that cells will not exit the region of interest from one frame to the next, and we exclude cells that are only partially visible in the current frames

**Fig. 2.**
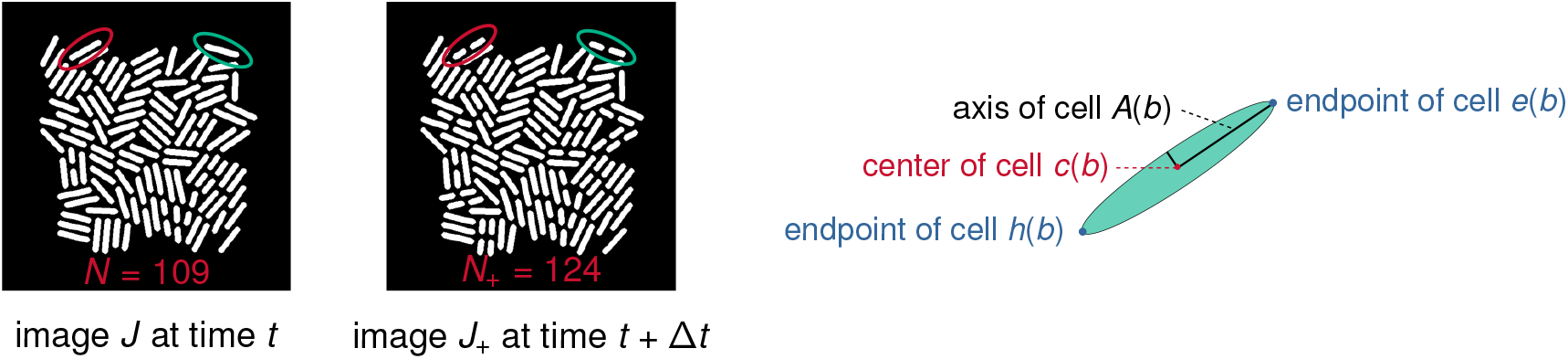
Simulated data and cell characteristics. Left: Two successive images generated by dynamic simulation for a colony of rod-shaped bacteria. Left image J displays N = 109 cells at time t. At time t + Δt with Δt = 1 min, cells have moved, grown, and some have divided. These cells are displayed in image J_+_, which contains N_+_ = 124 cells. We highlight two cells that have undergone a division between the frames (red and green ellipses). Right: Geometry of a rod shaped bacterium. Each cell b is identified by its center c(b), its long axis A(b), and the two endpoints e(b), h(b) of A(b).

## 3. Cell Characteristics

We next discuss characteristics of the *E. coli* bacteria important for our tracking algo- rithm.

### Cell Geometry

In accordance with the dynamics of bacterial colonies in microfluidic traps, the dynamic simulation software generates colonies of rod-shaped bacteria. Cell shapes can be approximated by long and thin ellipsoids, which are geometrically well identified by their center, their long axis, and the two endpoints of this long axis. The center *c*(*b*) is the centroid of all pixels belonging to cell *b*. The long axis *A*(*b*) of cell *b* is computed by principal component analysis (**PCA**). The endpoints *e*(*b*) and *h*(*b*) of cell *b* are the first and last cell pixels nearest to *A*(*b*); see Fig. 2 (right) for a schematic illustration.

### Cell Neighbors

For each image frame *J*, denote *B* = *B*(*J*) the set of fully visible cells in *J*, and by *N* = *N*(*J*) = card(*B*) the number of these cells. Let *V* be the set of all cell centers *c*(*b*) with *b* ∈ *B*. Denote *delV* the Delaunay triangulation [83] of the finite planar set *V* with *N* vertices. We say that two cells *b*_1_, *b*_2_ in *B* are *neighbors* if they verify the following three conditions:

1. (*b*_1_, *b*_2_) are connected by the edge *edg* of one triangle in *delV*.
2. The edge *edg* does not intersect any other cell in *B*.
3. Their centers verify ‖*c*(*b*_1_) − *c*(*b*_2_)‖ ≤*ρ*, where *ρ >* 0 is a user defined parameter.

For the synthetic images of size 600 × 600 that we considered (see Sec. 2), we take *ρ* = 80 pixels. We write *b*_1_∼ *b*_2_ for short, whenever *b*_1_, *b*_2_ are neighbors (i.e, satisfy the three conditions identified above).

### Cell Motion

Let *J, J*_+_ denote two successive images (i.e., frames). Denote *B* = *B*(*J*), *B*_+_ = *B*(*J*_+_) the associated sets of cells. Superpose temporarily the images *J* and *J*_+_ so that they then have the same center pixel. Any cell *b* ∈ *B*, which does not divide in the interframe *J* → *J*_+_, becomes a cell *b*_+_ in image *J*_+_. The “*motion vector*” of cell *b* from frame *J* to *J*_+_ is then defined by *v*(*b*) = *c*(*b*_+_) − *c*(*b*). If the cell *b* does divide between *J* and *J*_+_, denote *b*_div_ the last position reached by cell *b* at the time of cell division, and define similarly the motion *v*(*b*) = *c*(*b*_div_) − *c*(*b*). In our experimental recordings of real bacterial colonies with interframe duration 6 min, there is a *fixed number w >* 0 such that ‖*v*(*b*) ‖ ≤ *w/*2 for all cells *b*∈ *B*(*J*) for all pairs *J, J*_+_. In particular, we observed that for real image sequences, *w* = 100 pixels is an adequate choice. Consequently, we select *w* = 100 pixels for all simulated image sequences of BENCH6. For BENCH1 we select *w* = 45 pixels, again based on a comparison with real experimental recordings. Overall, the meta-parameter *w* is assumed to be a fixed number and to be known, since *w/*2 is an observable upper bound for the cell motion norm for a particular image sequence of a lab experiment.

### Target Window

Recall that *J, J*+ are temporarily superposed. Let *U* (*b*) ⊂ *J*_+_ be a square window of width *w*, with the same center as cell *b*. The *target window W* (*b*) is the set of all cells in *B*_+_ having their centers in *U* (*b*). Since ‖*v*(*b*) ‖ ≤ *w/*2, the cell *b*_+_ must belong to the target window *W* (*b*) ⊂ *B*_+_.

## 4. Registration Mappings

Next we discuss our assumptions on a valid registration mapping that establishes cell-to-cell correspondences between two frames. Let *J, J*_+_ denote two successive images, with cell sets *B* and *B*_+_, respectively. As above, we let *N* = card(*B*), and *N*_+_ = card(*B*_+_). Our goal is to track each cell from *J* to *J*_+_. For each cell *b* ∈ *B*, there exist three possible evolutions between *J* and *J*_+_:

**Case 1:** Cell *b* ∈ *B* did **not** divide in the interframe *J* →*J*_+_, and has become a cell *f* (*b*) ∈ *B*_+_; that is, *f* (*b*) has grown and moved during the interframe time interval.

**Case 2:** Cell *b* ∈ *B* divided between *J* and *J*_+_, and generated two children cells *b*_1_, *b*_2_ ∈ *B*_+_; we then denote *f* (*b*) = (*b*_1_, *b*_2_) ∈ *B*_+_ ×*B*_+_.

**Case 3:** Cell *b* ∈ *B* disappeared in the interframe *J* →*J*_+_, so that *f* (*b*) is not defined.

To simplify our exposition, we *ignore Case 3*, which can be handled by minor extensions of the cost minimization algorithms develop below. Consequently, a valid (true) registration mapping *f* will take values in the set {*B*_+_} ∪ {*B*_+_ × *B*_+_}.

## 5. Calibration of Cost Function Weights

With the notation we introduced, fix any two finite sets *A, A*_+_. Let *G* := {*g* : *A* → *A*_+_} be the set of all mappings *g* : *A* → *A*_+_. Fix *m* penalty functions pen_*k*_ (*g*) ≥ 0, *k* = 1, …, *m*. Let *g*^*^ ∈ *G* be the ground truth mapping we want to discover through minimization in *g* of some given cost function COST(*g*) defined by the linear combination of the penalty functions pen_*k*_ (*g*), the contributions of which are controlled by the cost function weights *λ*_*k*_ *>* 0. In this section, we present a generic *weight calibration algorithm*, extending a technique introduced and applied in [11, 12] for Markov random fields based image analysis.

The cost function must perform well (with the same weights) for hundreds of pairs of (synthetic) images *J, J*^+^. We consider one such synthetic pair for which the ground truth registration mapping *f* ∈ *G* is known, and use it to compute an adequate set of weights, which will then be used on all other synthetic pairs *J, J*^+^. Notice, that for experimental recordings of real cell colonies, no ground truth registration mappings *f* is available. In this case, *f* should be replaced by a set of user constructed, correct *partial* mappings defined on small subsets of *A*. The proposed weight calibration algorithm will also work in those situations.

We now show how knowing one ground truth mapping *f* can be used to derive the best feasible weights ensuring that *f* should be a plausible minimizer of the cost functional COST(*g*) over *g* ∈ *G*. Let PEN(*g*) = [pen_1_(*g*),…, pen_*m*_ (*g*)] be the vector of *m* penalties for any mapping *g* ∈ *G*. Let Λ = [*λ*_1_, …, *λ*_*m*_] be the weights vectors. Then, COST(*g*) = ⟨Λ, PEN(*g*) ⟩. Replacing *g* by another mapping *h* ≠ *g* induces the penalty changes ΔPEN_*g,h*_ = PEN(*h*) − PEN(*g*) and the cost change ΔCOST(*g, h*) = ⟨Λ, ΔPEN_*g,h*_ *⟩*. Now, fix any known ground truth mapping *f* ∈ *G*. We want *f* to be a minimizer of COST, so we should have

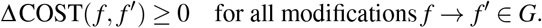

For each *a* ∈ *A*, select an arbitrary *s*(*a*) ∈ *W* (*a*) (where *W* (*a*) is the target window for cell *a*; see Sec. 3), to define a new mapping 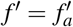 from *A* to *A*_+_ by 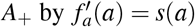, and 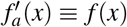 for all *x* ≠ *a*. Since *f* is a minimizer of COST, this single point modification 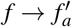 must generate the following cost increase

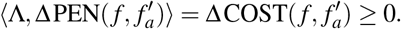

Denote *V*_*a*_ ∈ ℝ^*m*^ the vector 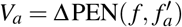. Then, the positive vector Λ ∈ ℝ ^*m*^, Λ ⪰ 0, should verify the set of linear constraints ⟨Λ,*V*_*a*_ ⟩ *≥*0 for all *a* ∈*A*. There may be too many such linear constraints. Consequently, we *relax* these constraints by introducing a vector *y* = [*y*(*a*)] ∈ ℝ ^card(*A*)^, *y ⪰*0, of slack variables *y*(*a*) ≥0 indexed by all the *a ∈ A*. (In optimization slack variables are introduced as additional unknowns to transform inequality constraints to an equality constraint and a non-negativity constraint on the slack variables.) We require the unknown positive vector Λ and the slack variables vector *y* to verify the system of linear constraints:

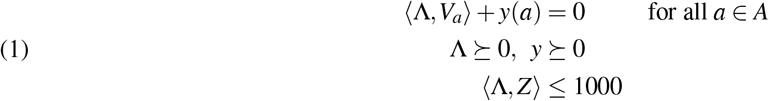

where *Z* = [1, …, 1] ∈ ℝ ^*m*^. The normalizing constant 1000 can be arbitrarily changed by rescaling. We seek high positive values for 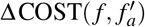 and small *L*_1_-norm for the slack variable vector *y*. So, we will seek two vectors Λ ∈ ℝ ^*m*^ and *y* ∈ ℝ ^card(*A*)^ solving the following *convex-concave* minimization problem, where *γ >* 0 is a user selected (large) meta parameter:

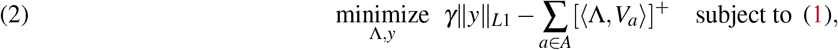

where we denote [*x*]^+^ := max(*x*, 0) for arbitrary *x*. To numerically solve the constrained minimization problem (2), we use the libraries `CVXPY` and `DCCP`(disciplined convex-concave programming) [2, 30, 80, 81]. DCCP is a package for convex-concave programming designed to solve non-convex problems. It can handle objective functions and constraints with any known curvature as defined by the rules of disciplined convex programming [24]. We give examples of numerically computed weight vectors Λ below. The computing time was less than 30 seconds for the data that we have prepared. For simplicity, we just considered one step changes in our computations, which make the overlap penalty weak. To increase the accuracy of the model it is possible to consider a larger number of samples (i.e., multi-step changes). Note that the solutions Λ of (2) are of course not unique, even after normalization by rescaling.

## 6. Cell Divisions and Children Pairing

In this section, we present the methodology for detecting cell divisions and our approach for pairing children with their parent cells.

### 6.1. Number *DIV*(*B, B*_+_) of Cell Divisions

Fix two successive synthetic image frames *J, J*_+_ with short inter-frame time equal to 1 minute. Their cell sets *B, B*_+_ have cardinality *N* and *N*_+_, respectively. We assume, for ease of presentation, that all cells *b* ∈ *B* still exist in *B*_+_, either as whole cells or divided into two children cells (i.e., no cells exit the field of view). This implies *N*_+_ ≥ *N*, and *DIV*(*B, B*_+_) = *N*_+_ −*N* is the number *DIV* of cell divisions occurring in the interframe *J* → *J*_+_.

Whenever *DIV >* 0, we want to compute the unknown set *trueCH* of true children pairs (*b*_1_, *b*_2_) ∈ *B*_+_ × *B*_+_. Each such pair is born from the division of some unknown parent cell *b* = parent(*b*_1_, *b*_2_). For the synthetic image sequence we should have card(*trueCH*) = *DIV*, but for computational advantage below, whenever *DIV*≥ 2 we *relax* this rigid constraint to the more pragmatic form |card(*trueCH*) −*DIV*| ≤ 1. For realistic experimental recordings, the relaxation bound is linked to the numbers of new cells entering *J*_+_ and of old cells exiting *J*_+_.

### 6.2. Linking Paired Children Cells to Parent Cells

For successive synthetic images *J, J*_+_ with 1 minute interframe such that *DIV*(*B, B*_+_) *>* 0, we call *PCH* the set of *plausible children pairs* defined as

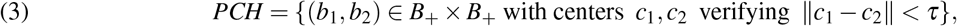

where the threshold *τ >* 0 is user selected and fixed for the whole benchmark set BENCH1 of synthetic image sequences.

To evaluate if a pair of cells (*b*_1_, *b*_2_)∈ *PCH* can qualify as a pair of children generated by division of a parent cell *b ∈ B*, we now quantify the “*geometric distortion*” between *b* and (*b*_1_, *b*_2_). Cell division of *b* into *b*_1_, *b*_2_ ∈ *B*_+_ occurs with small motion of *b*_1_, *b*_2_. During the short interframe duration the initial centers *c*_1_, *c*_2_ of *b*_1_, *b*_2_ in image *J* move by at most *w/*2 pixels each (see Sec. 3), and their initial distance to the center *c* of *b* is roughly at most ‖*A*(*b*) ‖ */*4, where *A*(*b*) is the long axis of cell *b*. Hence, the centers *c, c*_1_, *c*_2_ of *b, b*_1_, *b*_2_ should verify the constraint

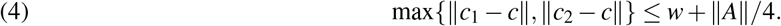

Define the set *SHLIN* of potential *short lineages* as the set all triplets (*b, b*_1_, *b*_2_) with *b* ∈ *B*, (*b*_1_, *b*_2_) ∈ *PCH*, verifying the preceding constraint (4). For each potential lineage (*b, b*_1_, *b*_2_) ∈*SHLIN*, define three terms penalizing the geometric distortions between a parent *b* ∈ *B* and a pair of children (*b*_1_, *b*_2_) ∈*PCH* by the following formulas, where we denote *c, c*_1_, *c*_2_ the centers of cells *b, b*_1_, *b*_2_ and *A, A*_1_, *A*_2_ their long axes

1. center distortion: cen(*b, b*_1_, *b*_2_) =‖ *c* − (*c*_1_ + *c*_2_)*/*2 ‖,
2. size distortion: siz(*b, b*_1_, *b*_2_) = |‖*A*‖ − (‖*A*_1_ ‖+ ‖*A*_2_ ‖) |,
3. angle distortion: ang(*b, b*_1_, *b*_2_) = angle(*A, A*_1_) + angle(*A, A*_2_) + angle(*A, c*_2_ − *c*_1_).

Here, angle denotes “*angles between non-oriented straight lines*,” and range from 0 to *π/*2.

Introduce three positive weights *λ*_cen_, *λ*_siz_, *λ*_ang_ (to be estimated), and for every short lineage (*b, b*_1_, *b*_2_)∈ *SHLIN* define its *distortion cost* by:

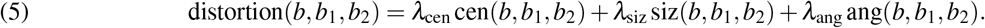

### 6.3. Estimating Most Likely Parent Cell for Children Pair

For each plausible pair of children (*b*_1_, *b*_2_) ∈ *PCH*, we will compute the most likely *parent cell b*^∗^ = parent(*b*_1_, *b*_2_) as the cell *b*^∗^ ∈ *B* minimizing distortion(*b, b*_1_, *b*_2_) in (5) over all *b* ∈ *B*, as summarized by the formula

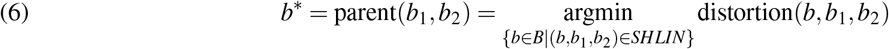

To force this minimization to yield a reliable estimate of *b*^∗^ = parent(*b*_1_, *b*_2_) for most true pairs of children (*b*_1_, *b*_2_), we calibrate the weights *λ*_*j*_, *j*∈{cen, siz, ang}by the algorithm outlined in section Sec. 5, using as “*ground truth set*” a fairly small set of visually identified true short lineages (*parent, children*).

For fixed (*b*_1_, *b*_2_), the set of potential parent cells *b*∈ *B* has very *small size* due to the constraint (4). Hence, brute force minimization of the functional distortion(*b, b*_1_, *b*_2_) in (5) over all *b* ∈*B* such that (*b, b*_1_, *b*_2_) ∈*SHLIN*, is a *fast computation* for each (*b*_1_, *b*_2_) in *PCH*. The distortion minimizing *b* = *b*^∗^ yields the most likely parent cell parent(*b*_1_, *b*_2_) = *b*^∗^.

### 6.4. Penalties to Control Children Pair Matching

True pairs of children cells *pch* = (*b*_1_, *b*_2_) ∈*PCH* must verify lineage and geometric constraints which we quantify via several penalties.

*“Lineage” Penalty*.. Valid children pairs (*b*_1_, *b*_2_) ∈*PCH* should be correctly matchable with the their most likely parent cell *b*^∗^ = parent(*b*_1_, *b*_2_) (see (6)). So, we define the *“lineage” penalty* lin(*b*_1_, *b*_2_) = distortion(*b*^∗^, *b*_1_, *b*_2_) by

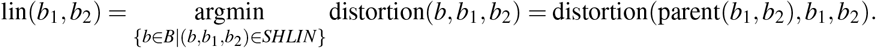

Notice that the computation of lin(*b*_1_, *b*_2_) is quite fast.

*“Gap” Penalty*.. Denote *tips*(*b*) the set of two endpoints of any cell *b*. For any pair *pch* = (*b*_1_, *b*_2_) ∈ *PCH*, define endpoints *x*_1_ ∈ *tips*(*b*_1_), *x*_2_ ∈ *tips*(*b*_2_) and the *gap* penalty gap(*b*_1_, *b*_2_) by

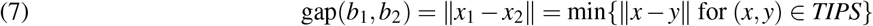

with *TIPS* = *tips*(*b*_1_) × *tips*(*b*_2_).

*“Dev” Penalty*.. For rod-shaped bacteria, a true pair (*b*_1_, *b*_2_) ∈*PCH* of just born children must have a small gap(*b*_1_, *b*_2_) = ‖*x*_1_ − *x*_2_ ‖and roughly aligned cells *b*_1_ and *b*_2_. For (*b*_1_, *b*_2_) ∈*PCH*, we quantify the *deviation from alignment* dev(*b*_1_, *b*_2_) as follows. Let *x*_1_, *x*_2_ be the closest endpoints of *b*_1_, *b*_2_ (see (7)). Let *str*_12_ be the straight line linking the centers *c*_1_, *c*_2_ of *b*_1_, *b*_2_. Let *d*_1_, *d*_2_ be the distances from *x*_1_, *x*_2_ to the line *str*_12_. Set then

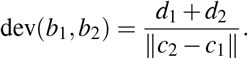

*“Ratio” Penalty*.. True children pairs must have nearly equal lengths. So, for (*b*_1_, *b*_2_) ∈ *PCH* with lengths *L*_1_, *L*_2_ we define the length *ratio penalty* by

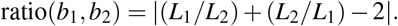

*“Rank” penalty*.. Let *L*_min_ be the minimum cell length over all cells in *B*_+_. In *B*_+_, children pairs (*b*_1_, *b*_2_) just born during interframe *J* → *J*_+_ must have lengths *L*_1_, *L*_2_ close to *L*_min_. So, for (*b*_1_, *b*_2_) ∈ *PCH*, we define the *rank* penalty by

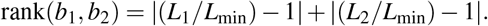

### 6.5. Cost Function Dedicated to Children Pairing

Given two successive images *J, J*_+_ with a positive number *DIV* = *N*_+_(*J*) − *N*(*J*) of cell divisions, the set *trueCH* ⊂*B*_+_ × *B*_+_ of true children pairs will have size *DIV* and be a subset of the set *PCH* of plausible children pairs. In the search for *trueCH*, the unknown is a subset *X* of *PCH*. We build a cost function *ϕ* (*X*) approximately minimized when *X* is close to *trueCH*. The main term of *ϕ* (*X*) will be the sum over all pairs in *X* of a weighted linear combination of the penalty functions {lin, gap, dev, ratio, rank}. Another penalty will ensure that *X* contains no overlapping pairs. The minimization of *ϕ* (*X*), as well as of several further combinatorial cost functions below will be implemented by intensive simulations of BMs. We present these stochastic neural networks, next.

## 7. Generic Boltzmann Machines (BMs)

Minimization of our main cost functionals is a heavily combinatorial task, since the unknown variable is a mapping between two finite sets of sizes ranging from 80 to 120. To handle these minimizations, we use BMs originally introduced by Hinton et al. (see [1, 38]). Indeed, these recurrent stochastic neural networks can efficiently emulate some forms of simulated annealing.

Each BM implemented here is a network *BM* = {*U*_1_, …, *U*_*N*_}of *N stochastic neurons U*_*j*_. At time *t* = 0, 1, 2, …, each neuron *U*_*j*_ has a random state *Z*_*j*_(*t*) belonging to a fixed finite set *W* (*j*). The *configuration Z*(*t*) = {*Z*_1_(*t*), …, *Z*_*N*_(*t*)}of the whole network *BM* thus belongs to the *configurations set CONF* = *W* (1) ×…×*W* (*N*). Neurons interactivity is specified by a finite set *CLQ* of *cliques*. Each clique *K* is a subset of *S* = {1, …, *N*}. During configuration updates *Z*(*t*) → *Z*(*t* + 1), neurons may interact only if they are in the same clique. Here, all cliques *K* are of small sizes 1, or 2, or 3.

For each clique *K*, one specifies an energy function *J*_*K*_(*z*) defined for all *z* ∈ *CONF*, with *J*_*K*_(*z*) depending only on the *z* _*j*_ such that *j* ∈ *K*. The full energy *E*(*z*) of configuration *z* is then defined by

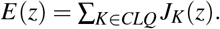

The BM stochastic dynamics *Z*(*t*) → *Z*(*t* + 1) is driven by the energy function *E*(*z*), and by a fixed decreasing sequence of *virtual temperatures Temp*(*t*) *>* 0, tending slowly to 0 as *t* → ∞. Here we use standard temperature schemes of the form *Temp*(*t*) ≡ *cη*^*t*^ with fixed *c >* 0 and slow *decay rate* 0.99 *< η <* 1.

We have implemented the classical “*asynchronous*” BM dynamics. At each time *t, only one* random neuron *U*_*j*_ may modify its state, after reading the states of all neurons belonging to cliques containing *U*_*j*_. A much faster alternative, implementable on GPUs, is the “*synchronous*” BM dynamics, where at each time *t* roughly 50% of all neurons may simultaneously modify their states (see [10, 9, 14]). The detailed BM dynamics is presented in the appendix (see Sec. A).

Recall that when the virtual temperatures *Temp*(*t*) decrease slowly enough to 0, the energy *E*(*Z*(*t*)) converges in probability to a local minimum of the BM energy *E*(*z*) over all configurations *z* ∈ *CONF*.

## 8. Optimized Children Pairing

Next, we present a formulation of an optimization problem to pair parents with their children. Fix successive images *J, J*_+_ with a positive number of cell divisions *DIV* = *N*_+_ − *N*. Denote *PCH* = {*pch*_1_, *pch*_2_,…, *pch*_*m*_} the set of *m* plausible children pairs (*b*_1_, *b*_2_) in *B*_+_. The penalties lin, gap, dev, ratio, and rank defined above for all pairs (*b*_1_, *b*_2_) ∈ *PCH* determine five numerical vectors *LIN, GAP, DEV, RAT, RANK* in ℝ^*m*^ with coordinates *LIN*_*j*_ = lin(*pch*_*j*_), *GAP*_*j*_ = gap(*pch*_*j*_), *DEV*_*j*_ = dev(*pch*_*j*_), *RAT*_*j*_ = ratio(*pch*_*j*_), *RANK*_*j*_ = rank(*pch*_*j*_).

We now define a *binary* BM constituted by *m binary* stochastic neurons *U*_*j*_, *j* = 1 … *m*. At time *t* = 0, 1, 2, …, each *U*_*j*_ has a random *binary valued state Z*_*j*_(*t*) = 1 or 0. The random configuration *Z*(*t*) = [*Z*_1_(*t*), …, *Z*_*m*_(*t*)] of this BM belongs to the configuration space *CONF* = {0, 1} ^*m*^ of all binary vectors *z* = [*z*_1_, …, *z*_*m*_]. Let *SUB* bet the set of all subsets of *PCH*. Each configuration *z* ∈ *CONF* is the indicator function of a subset *sub*(*z*) of *PCH*. We view each *sub*(*z*) ∈ *SUB* as a possible estimate for the unknown set *trueCH* ⊂ *B*_+_ × *B*_+_ of true children pairs (*b*_1_, *b*_2_). For each potential estimate *sub*(*z*) of *trueCH*, the “*lack of quality*” of the estimate *sub*(*z*) will be penalized by the *energy function E*(*z*) ≥ 0 of our binary BM. We now specify the energy *E*(*z*) for all *z* ∈ *CONF* by combining the penalty terms introduced above.

True children pairs born from distinct parents must clearly not intersect. To enforce this constraint, define the symmetric *m* × *m* binary matrix [*Q*_*j,k*_] by **i)** *Q*_*j,k*_ = 1 if *j* ≠ *k* and the two pairs *pch*_*j*_, *pch*_*k*_ have **one** cell in common, **ii)** *Q*_*j,k*_ = 0 if *j* ≠ *k* and the two pairs *pch*_*j*_, *pch*_*k*_ have **no** cell in common, **iii)** *Q*_*j, j*_ = 0 for all *j*.

The quadratic penalty *z ↦ ⟨z, Qz⟩* is non-negative for *z* ∈ *CONF*, and must be zero if *sub*(*z*) = *trueCH*. Introduce six positive weight parameters to be selected further on *λ*_*j*_, *j*∈ {lin, gap, dev, rat, rank, *Q*}. Define the vector *V*∈ ℝ^*m*^ as a weighted linear combination of the penalty vectors *LIN, GAP, DEV, RAT, RANK*

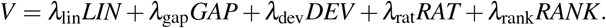

For any configuration *z* ∈ *CONF*, the BM energy *E*(*z*) is defined by the *quadratic function*

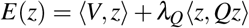

We already know that the unknown set *trueCH* of true children pairs must have cardinal *DIV* = *N*_+_ − *N*. So we seek a configuration *z*^∗^ ∈ *CONF* minimizing the energy *E*(*z*) under the rigid constraint card {*sub*(*z*)} = *DIV*. Let *ONE* ∈ ℝ^*m*^ be the vector with all its coordinates equal to 1. The constraint on *z* can be reformulated as ⟨*ONE, z⟩* = *DIV*. We want the unknown *trueCH* to be close to the solution *z*^∗^ of the constrained minimization problem

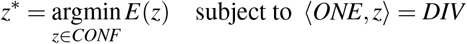

To force this minimization to yield a reliable estimate of *trueCH*, we calibrate the six weights

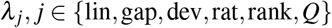

by the algorithm in Sec. 5. Denote *CONF*_1_ the set of all *z* ∈ *CONF* such that ⟨*ONE, z ⟩*= *DIV*. To minimize *E*(*z*) under the constraint *z* ∈ *CONF*_1_, fix a slowly decreasing temperature scheme *Temp*(*t*) as in Sec. 7. We need to force the BM stochastic configurations *Z*(*t*) to remain in *CONF*_1_. Then, for large time step *t*, the *Z*(*t*) will converge in probability to a configuration *z*^∗^ ∈ *CONF*_1_ approximately minimizing *E*(*z*) under the constraint *z* ∈ *CONF*_1_.

Start with any *Z*(0) ∈ *CONF*_1_. Assume that for 0 ≤ *s* ≤ *t*, one has already dynamically generated BM configurations *Z*(*s*) ∈ *CONF*_1_. Then, randomly select two sites *j, k* such that *Z*_*j*_(*t*) = 1 and *Z*_*k*_(*t*) = 0. Compute a virtual configuration *Y* by setting *Y*_*j*_ = 0, *Y*_*k*_ = 1, and *Y*_*i*_ ≡ *Z*_*i*_ for all sites *i* different from *j* and *k*. Compute the energy change Δ*E* = *E*(*Y*) − *E*(*Z*(*t*)), and the probability *p*(*t*) = exp(−*D/Temp*(*t*)), where *D* = max{0, Δ*E*}. Then randomly select *Z*(*t* + 1) = *Y* or *Z*(*t* + 1) = *Z*(*t*) with respective probabilities *p*(*t*) and (1 − *p*(*t*)). Clearly, this forces *Z*(*t* + 1) ∈ *CONF*_1_

## 9. Performance of Automatic Children Pairing on Synthetic Videos

In the following subsections we provide experimental results for pairing children and parent cells.

### 9.1. Children Pairing: Fast BM simulations

For *m* = card(*PCH*) ≤ 1000, one can reduce the computational cost for BM dynamics simulations by pre-computing and storing the *m* × *m* symmetric binary matrix *Q*, as well as the *m*-dimensional vectors *LIN, GAP, DEV, RAT, RANK* and their linear combination *V*. A priori reduction of *m* significantly reduces the computing times, and can be implemented by trimming away the pairs *pch*_*j*_ ∈ *PCH* for which the penalties *LIN* _*j*_, *GAP*_*j*_, *DEV* _*j*_, *RAT* _*j*_, and *RANK* _*j*_ are all larger than predetermined empirical thresholds. We performed a study on 100 successive (synthetic) images. We show scatter plots for the most informative penalty terms in Fig. 3. These plots allow us to determine adequate thresholds for the penalty terms. We observed that for the synthetic and real data we considered the trimming of *DEV, GAP*, and *RANK* reduced the percentage of invalid children pairs by 95%, therefore drastically reducing the combinatorial complexity of the problem.

**Fig. 3.**
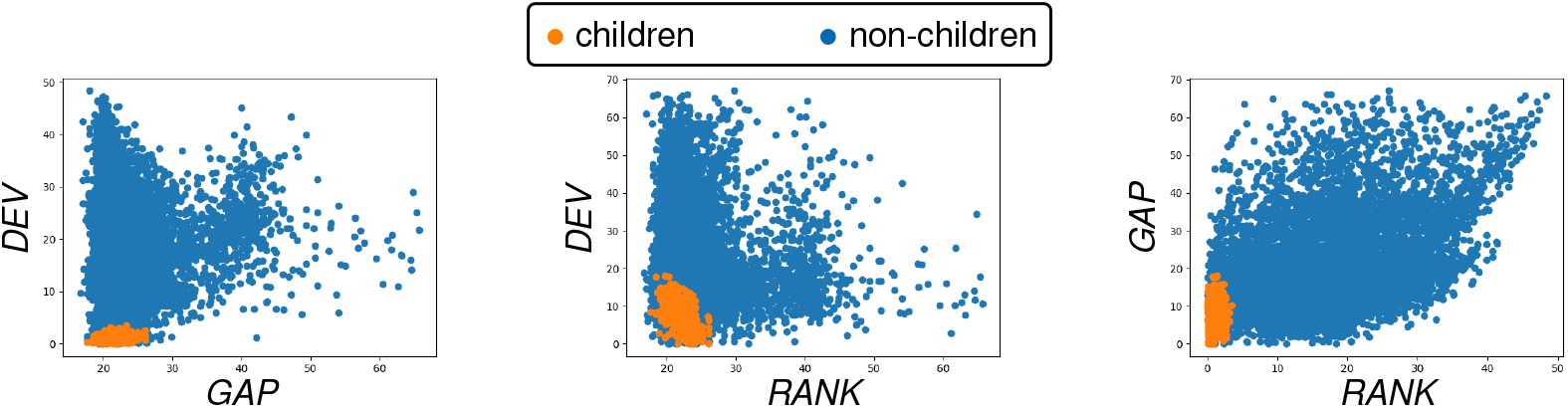
Scatter plots for tandems of the penalty terms DEV, GAP, and RANK. We mark in orange the true children pairs and in blue invalid children pairs. These plots allow us to identify appropriate empirical thresholds to trim the (considered synthetic) data in order to reduce the computational complexity of the parent-children pairing.

The quadratic energy function *E*(*z*) is the sum of clique energies *J*_*K*_(*z*) involving only cliques of cardinality 1 and 2. For any clique *K* ={*j*} of cardinality 1, with 1 ≤*j* ≤*m*, one has *J*_*K*_(*z*) = *V*_*j*_*z* _*j*_. For any clique *K* ={*j, k*}of cardinality 2, with 1≤ *j < k* ≤*m*, one has *J*_*K*_(*z*) = 2*Q*_*j,k*_*z* _*j*_*z*_*k*_. A key computational step when generating *Z*(*t* + 1) is to evaluate the energy change Δ*E* when one flips the binary values *Z*_*j*_(*t*) = 1 and *Z*_*k*_(*t*) = 0 by the new value (1− *z*_*i*_) for a fixed single site *i*. This step is quite fast since it uses only the numbers *V*_*j*_, *V*_*k*_, and ⟨*q*(*j*), *Z*(*t*) ⟩, ⟨*q*(*k*), *Z*(*t*) ⟩, where *q*(*i*) is the *i*^*th*^ row of the matrix *Q*.

### 9.2. Children Pairing: Implementation on Synthetic Videos

We have implemented our children pairing algorithms on synthetic image sequences having 100 to 500 image frames with 1 minutes interframe (benchmark set BENCH1; see Sec. 2). The cell motion bound *w/*2 per interframe was defined by *w* = 20 pixels. The parameter *τ* that defines the sets *PCH* of plausible children pairs (see (3)) was set at *τ* = 45 pixels.

The known true cell registrations indicated that in our typical BENCH1 image sequence, the successive sets *PCH* had average cardinals of 120, while the number of true children pairs per *PCH* roughly ranged from 2 to 6 with a median of 4. The size of the reduced configuration space *CONF*1 per image frame thus ranged from 10^4^ to 120^6^*/*6! = 4.2 ·10^9^ with a median of 9· 10^6^

Our weights estimation technique introduced in Sec. 5 yields the weights [*λ*_cen_, *λ*_siz_, *λ*_ang_] = [0.255, 0.05, 0.05] and [*λ*_gap_, *λ*_dev_, *λ*_rat_, *λ*_rank_] = [0.01, 1, 0.0001, 0.05] or the penalties introduced in Sec. 13. To reduce the computing time for hundreds of BM energy minimizations on the BENCH1 image sequences, we excluded obviously invalid children pairs in each *PCH* set, by simultaneously thresholding of the penalty terms. The BM temperature scheme was *Temp*(*t*) = 1000 (0.995)^*t*^, with the number of epochs capped at 5000. The average CPU time for BM energy minimization dedicated to optimized children pairing was about 30 seconds per frame.

### 9.3. Parent-Children Matching: Accuracy on Synthetic Videos

For each successive image pair *J, J*_+_, with cells *B, B*_+_ of cardinality *N < N*_+_, our parent-children matching algorithm computes a set *SHL* of short lineages (*b, b*_1_, *b*_2_), where the cell *b*∈ *B* is expected to be the parent of cells *b*_1_, *b*_2_ ∈*B*_+_. Recall that *DIV* = *N*_+_ − *N* provides the number of cell divisions during the interframe *J* →*J*_+_. The number *VAL* of correctly reconstructed short lineages (*b, b*_1_, *b*_2_) ∈*SHL* is obtained by direct comparison to the known ground truth registration *J*→ *J*_+_. For each frame *J*, we define the *pcp-accuracy* of our Parent-Children Pairing algorithm as the ratio *VAL/DIV*.

We have tested our parent-children matching algorithm on three long synthetic image sequences BENCH1 (500 frames), BENCH2 (300 frames), and BENCH3 (300 frames), with respective interframes of 1, 2, and 3 minutes. For each frame *J*_*k*_, we computed the pcp-accuracy between *J*_*k*_ and *J*_*k*+1_.

We report the accuracies of our parent-children pairing algorithms in Tab. 1. For BENCH1, all 500 pcp-accuracies reached 100%. For BENCH2, pcp-accuracies reach 100% for 298 frames out of 300, and for the remaining two frames, accuracies were still high at 93% and 96%. For BENCH3, where interframe duration was longest (3 minutes), the 300 pcp-accuracies decreased slightly but still averaged 99%, and never fell below 90%.

**Table 1.**
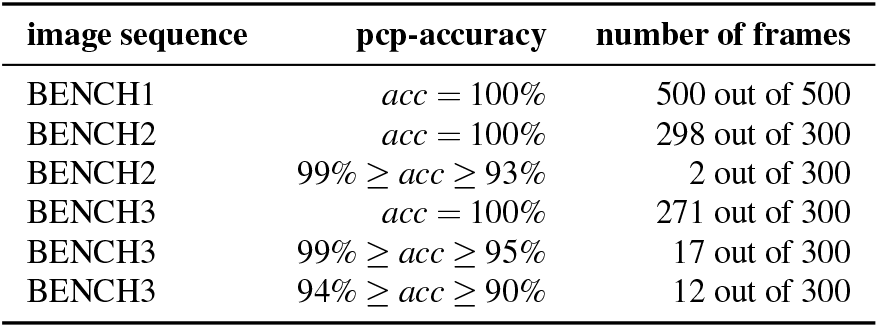
Accuracies of parent-children pairing algorithm. We applied our parent-children pairing algorithm to three long synthetic image sequences BENCH1 (500 frames), BENCH2 (300 frames), and BENCH3 (300 frames), with interframe intervals of 1, 2, 3 minutes, respectively. The table summarizes the resulting pcp-accuracies. Note that pcp-accuracies are practically always at 100%. For BENCH2 pcp-accuracies are 100% for 298 frames out of 300, and for the remaining two frames, accuracies were still high at 93% and 96%. For BENCH3 the average pcp-accuracy for the 3 minute interframe is 99%.

## 10. Reduction to Registrations with No Cell Division

Fix successive frames *J, J*_+_ and their cell sets *B, B*_+_. We seek the unknown registration mapping *f* : *B* → {*B*_+_ ∪ (*B*_+_ × *B*_+_)}, where *f* (*b*) ∈ *B*_+_ iff cell *b* did not divide during the interframe *J* → *J*_+_ and *f* (*b*) = (*b*_1_, *b*_2_) ∈ *B*_+_ × *B*_+_ iff cell *b* divided into (*b*_1_, *b*_2_) during the interframe.

If card(*B*) = *N < N*_+_ = card(*B*_+_), we know that the number of cell divisions during the interframe *J* → *J*_+_ should be *DIV* = *DIV*(*B, B*_+_) = *N*_+_ − *N >* 0. We then apply the parent-children matching algorithms outlined above to compute a set *SHL* = *SHL*(*B, B*_+_) of short lineages (*b, b*_1_, *b*_2_) with *b* ∈ *B, b*_1_, *b*_2_ ∈ *B*_+_ and card(*SHL*) = *DIV*. For each (*b, b*_1_, *b*_2_) ∈ *SHL*, the cell *b* is computed by *b* = parent(*b*_1_, *b*_2_) as the parent cell of the two children cells *b*_1_, *b*_2_ ∈ *B*_+_.

For each (*b, b*_1_, *b*_2_) ∈ *SHL*, eliminate from *B* the parent cell, *b*, and eliminate from *B*_+_ the two children cells *b*_1_, *b*_2_. We are left with two residual sets, *resB* ⊂ *B* and *resB*_+_ ⊂ *B*_+_, having the same cardinality, *N* − *DIV* = *N*_+_ − 2*DIV*. Assuming that our set *SHC* of short lineages is correctly determined, the cells *b*∈ *redB* should not divide in the interframe *J* → *J*_+_, and hence have a single (still unknown) registration *f* (*b*) ∈ *redB*_+_. Thus, the still unknown part of the registration *f* is a bijection from *redB* to *redB*_+_.

Let *divB* = *B* − *redB* and *divB*_+_ = *B*_+_ − *redB*_+_. For each *b* ∈ *divB*, the cell *b* divides into the unique pair of cells, (*b*_1_, *b*_2_) ∈ *divB*_+_ × *divB*_+_, such that (*b, b*_1_, *b*_2_) ∈ *SHL*. Hence, we can set *f* (*b*) = (*b*_1_, *b*_2_) for all *b* ∈ *divB*. Thus, the remaining problem to solve is to compute the bijective registration *f* : *redB*→ *redB*_+_. We have reduced the registration discovery to a new problem, where *no cell divisions occur* in the interframe duration. In what follows, we present our algorithm to solve this registration problem.

## 11. Automatic Cell Registration after Reduction to Cases with No Cell Division

As indicated above, we can *explicitly reduce* the generic cell tracking problem to a problem where there is *no cell division*. We consider images *J, J*_+_ with associated cell sets *B, B*_+_ such that *N* = card(*B*) = card(*B*_+_). Hence, there are no cell divisions in the interframe *J* →*J*_+_ and the map *f* of this reduced problem is (in principle) a bijection *f* : *B*→ *B*_+_ with card(*B*) = card(*B*_+_). We show two typical successive images we use for testing with no cell division generated by the simulation software [94] (see Sec. 2) in Fig. 4.

**Fig. 4.**
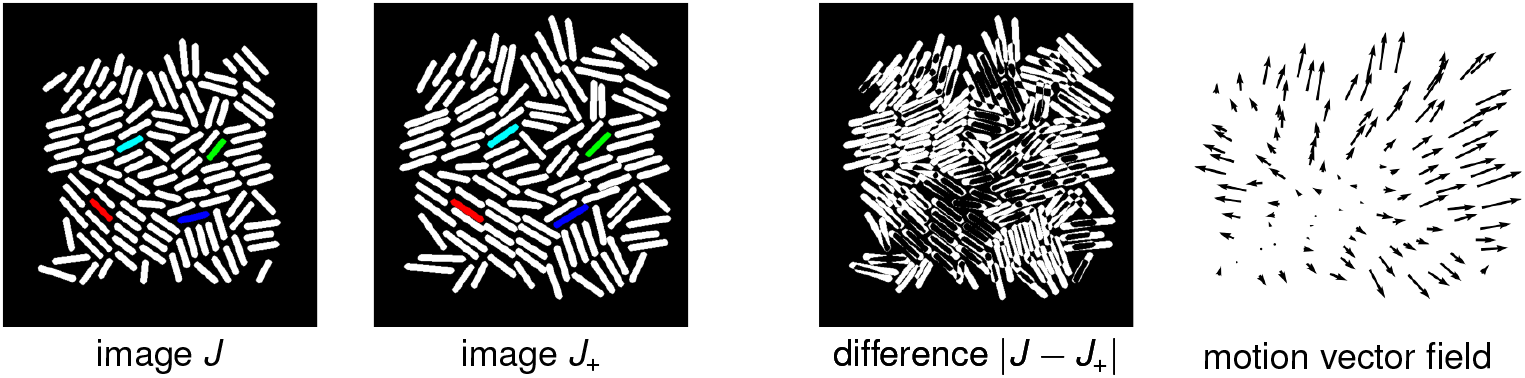
Simulated cell dynamics. From left to right, two successive simulated images J and J_+_ with an interframe time of six minutes and no cell division, their image difference| J − J_+_ |, and the associated motion vectors. For the image J and J_+_ we color four pairs of cells in B × B_+_, which should be matched by the true cell registration mapping. Notice that the motion for an interframe time of six minutes is significant. We can observe that even without considering cell division, we can no longer assume that corresponding cells in frame J and J_+_ overlap.

### 11.1. The set *MAP* of Many-to-One Cell Registrations

We have reduced the registration search to a situation where during the interframe *J* → *J*_+_, no cell has divided, no cell has disappeared, and no cell has suddenly emerged in *B*_+_ without originating from *B*. The unknown registration *f* : *B*→ *B*_+_ should then in principle be injective and onto. However, for computational efficiency, we will temporarily relax the bijectivity constraint on *f*. We will seek *f* in the set *MAP* of all *many-to-one mappings f* : *B* → *B*_+_ such that for each *b* ∈ *B*, the cell *f* (*b*) is in the target window *W* (*b*) ⊂ *B*_+_ (see section Sec. 3).

### 11.2. Registration Cost Functional

To design a cost functional cost(*f*), which should be roughly minimized when *f* ∈ *MAP* is very close to the true registration from *B* to *B*_+_, we linearly combine penalties match(*f*), over(*f*), stab(*f*), flip(*f*) weighted by unknown positive weights *λ*_match_, *λ*_over_, *λ*_stab_, *λ*_flip_, to write, for all registrations *f* ∈ *MAP*,

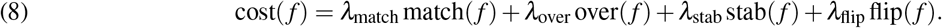

We specify the individual terms that appear in (8) below. Ideally, the minimizer of cost(*f*) over all *f* ∈ *MAP* is close to the unknown true registration mapping *f* : *B* → *B*_+_. To enforce a good approximation of this situation, we first estimate efficient positive weights by applying our calibration algorithm (see Sec. 5). The actual minimization of cost(*f*) over all *f* ∈ *MAP* is then implemented by a BM described in Sec. 12.

#### 11.2.1. Cell Matching Likelihood

match(*f*). Here, we extend a pseudo likelihood approach used to estimate parameters in Markov random fields modeling by Gibbs distributions (see [49]). Recall that *g*.*rate* is the *known* average cell growth rate. For any cells *b* ∈ *B, b*_+_ ∈ *B*_+_, the geometric quality of the matching *b ↦b*_+_ relies on three main characteristics: (*i*) motion *c*(*b*_+_) − *c*(*b*) of the cell center *c*(*b*), (*ii*) angle between the long axes *A*(*b*) and *A*(*b*_+_), (*iii*) cell length ratio ‖*A*(*b*_+_)‖ */ ‖A*(*b*) ‖. So, for all *b*∈ *B* and *b*_+_ in the target window *W* (*b*), define

1. Kinetic energy: kin(*b, b*_+_) = ‖*c*(*b*) − *c*(*b*_+_)‖ ^2^.
2. Distortion of cell length:

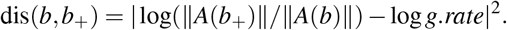
3. Rotation angle: 0 ≤ rot(*b, b*_+_)≤ *π/*2 is the geometric angle between the straight lines carrying *A*(*b*) and *A*(*b*_+_).

Fix *b* ∈ *B*, and let *b′* run through the whole target window *W* (*b*). The finite set of values thus reached by the kinetic penalties kin(*b, b′*) has two smallest values *kin*_1_(*b*), *kin*_2_(*b*). Define *list*.*kin* =⋃ _*b* ∈ *B*_ {kin_1_(*b*), kin_2_(*b*)}, which is a list of 2*N* “*low*” kinetic penalty values. Repeat this procedure for the penalties dis(*b, b′*) and rot(*b, b′*) to similarly define a *list*.*dis* of 2*N* “*low*” distortion penalty values, and a *list*.*rot* of 2*N* “*low*” rotation penalty values.

The three sets *list*.*kin, list*.*dis, list*.*rot* can be viewed as three random samples of size 2*N*, respectively, generated by three unknown probability distributions *P*_kin_, *P*_dis_, *P*_rot_. We approximate these three probabilities by their *empirical* cumulative distribution functions CDF_kin_, CDF_dis_, CDF_rot_, which can be readily computed. We now use the right tails of these three CDFs to compute separate probabilistic evaluations of how *likely* the matching of cell *b* ∈ *B* with cell *b*_+_ ∈ *W* (*b*) is. For any fixed mapping *f* ∈ *MAP*, and any *b* ∈ *B*, set *b*_+_ = *f* (*b*). Compute the three penalties *vkin* = kin(*b, b*_+_), *vdis* = dis(*b, b*_+_), *vrot* = rot(*b, b*_+_), and define three associated “*likelihoods*” for the matching *b* → *b*_+_ = *f* (*b*).

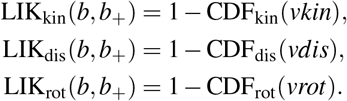

High values of the penalties *vkin, vdis, vrot* thus will yield three small likelihoods for the matching *b* → *b*_+_ = *f* (*b*). With this, we can define a “*joint likelihood*” 0 ≤ LIK(*b, b*_+_) ≤ 1 evaluating how likely is the matching *b* → *b*_+_ = *f* (*b*):

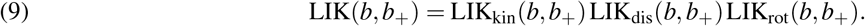

Note that higher values of LIK(*b, b*_+_) correspond to a better geometric quality for the matching of *b* with *b*_+_ = *f* (*b*). To avoid vanishingly small likelihoods, whenever LIK(*b, b*_+_) *<* 1e−6, we replace it by 1e−6. Then, for any mapping *f* ∈ *MAP*, we define its *likelihood* lik(*f*) by the finite product

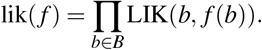

The product of these *N* likelihoods is typically very small, since *N* = card(*B*) can be large. So, we evaluate the geometric matching quality match(*f*) of the mapping *f* via the averaged *log-likelihood of f*, namely,

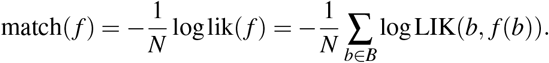

Good registrations *f* ∈ *MAP* should yield small values for the criterion match(*f*).

#### 11.2.2. Overlap

over(*f*) We expect *bona fide* cell registrations *f* ∈ *MAP* to be bijections. Consequently, we want to penalize mappings *f* which are many-to-one. We say that two distinct cells (*b, b′*) ∈ *B* × *B* do *overlap* for the mapping *f* ∈ *MAP* if *f* (*b*) = *f* (*b′*). The total number of overlapping pairs (*b, b′*) for *f* defines the *overlap penalty*:

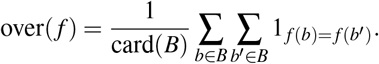

#### 11.2.3. Neighbor Stability

stab(*f*) Let *B* = {*b*_1_, …, *b*_*N*_}. Denote *G*_*i*_ the set of all neighbors for cell *b*_*i*_ in *B* (i.e., *b* _*j*_ ∼*b*_*i*_ ⇔*b* _*j*_ ∈ *G*_*i*_; see Sec. 3). For *bona fide* registrations *f* ∈ *MAP*, and for most pairs of neighbors *b*_*i*_ ∼ *b* _*j*_ in *B*, we expect *f* (*b*_*i*_) and *f* (*b*_*j*_) to remain neighbors in *B*_+_. Consequently, we penalize the lack of “*neighbors stability*” for *f* by

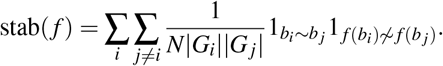

#### 11.2.4. Neighbor Flip

flip(*f*) Fix any mapping *f* ∈ *MAP*, any cell *b* ∈ *B* and any two neighbors *b′, b″* of *b* in *B*. Let *z* = *f* (*b*), *z′* = *f* (*b′*), *z″* = *f* (*b ″*). Let *c, c′, c″* and *d, d ′, d″* be the centers of cells *b, b′, b″* and *z, z′, z″*. Let *α* be the oriented angle between *c ′− c* and *c″* − *c*, and let *α*_*f*_ be the angle between *d ′− d* and *d″ − d*, respectively. We say that the mapping *f* has flipped cells *b′, b″* around b, and we set FLIP(*f, b, b′, b″*) = 1 if *z′, z″* are both neighbors of *z*, and the two angles *α, α*_*f*_ have *opposite signs*. In all other cases, we set FLIP(*f, b, b′, b ″*) = 0.

For any registration *f* ∈ *MAP*, define the *flip penalty* for *f* by

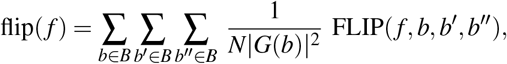

where *G*(*b*) is the neighborhood of cell *b* in *B*. In Fig. 5 we illustrate an example of an unwanted cell flip.

**Fig. 5.**
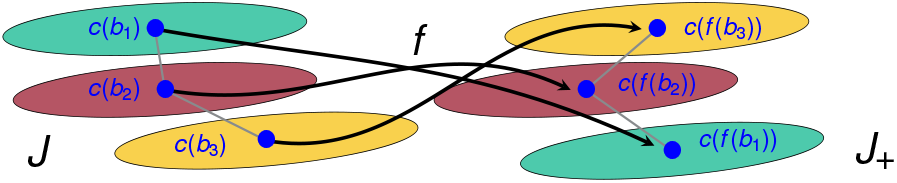
Illustration of an undesirable flip for the mapping f. The cells b_1_ and b3 are neighbors of b_2_, and mapped by f on neighbors z1 = f (b_1_), z3 = f (b3) of z2 = f (b_2_), as should be expected for bona fide cells registrations. But for this mapping f, we have z3 above z2 above z1, whereas for the original cells we had b_1_ above b_2_ above b3. Our cost function penalizes flips of this nature.

## 12. BM Minimization of Registration Cost Function

In what follows, we define the optimization problem for the registration of cells from one frame to another (i.e., cell tracking), as well as associated methodology and parameter estimates.

### 12.1. BM Minimization of cost(*f*) over *f* ∈ *MAP*

Let *B, B*_+_ be two successive sets of cells. As outlined above, we have reduced the problem to one in which we can assume that *N* = card(*B*) = card(*B*_+_), so that there is no cell division during the interframe. Write *B* ={*b*_1_, …, *b*_*N*_}. For short, denote *W* (*j*) ⊂ *B*_+_ instead of *W* (*b*_*j*_) the target window of cell *b* _*j*_. We seek to minimize cost(*f*) over all registrations *f* ∈ *MAP*. Let *BM* be a BM with sites *S* = {1, …, *N*}and stochastic neurons {*U*_1_, …, *U*_*N*_}. At time *t*, the random state *Z*_*j*_(*t*) of *U*_*j*_ will be some cell *z* _*j*_ belonging to the target window *W* (*j*) and the random configuration *Z*(*t*) = {*Z*_1_(*t*), …, *Z*_*N*_(*t*)}of the whole *BM* belongs to the configurations set *CONF* = *W* (1) × … ×*W* (*N*).

To any configuration *z* = {*z*_1_, …, *z*_*N*_}∈ *CONF*, we associate a unique cell registration *f* ∈*MAP* defined by *f* (*b*_*j*_) = *z* _*j*_ for all *j*, denoted by *f* = map(*z*). This determines a bijection *z↦ f* = map(*z*) from *CONF* onto *MAP*. The inverse of map : *CONF* → *MAP* will be called range : *MAP* →*CONF*, and is defined by *z* = range(*f*), when *z* _*j*_ = *f* (*b*_*j*_) for all *j*.

### 12.2. BM Energy Function *E*(*z*)

We now define the energy function *E*(*z*) ≥ 0 of our BM for all *z* ∈ *CONF*. Denote *E*^∗^ = minimize_*z*∈*CONF*_ *E*(*z*). Since *f* ↦ *z* = range(*f*) is a bijection from *MAP* to *CONF*, we must have

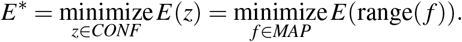

Our goal is to minimize cost(*f*), and we know that BM simulations should roughly minimize *E*(*z*) over all *z* ∈ *CONF*. So, we define the BM energy function *E*(*z*) by forcing

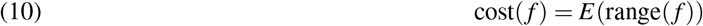

for any registration mapping *f* ∈ *MAP*, which—due to the preceding subsection—is equivalent to

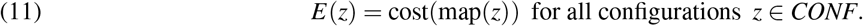

The next subsection will explicitly express the energy *E*(*z*) in terms of *cliques* of neurons. Due to (10) and (11) we have

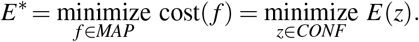

For large time *t*, the BM stochastic configuration *Z*(*t*) tends with high probability to concentrate on configurations *z* ∈*CONF*, which roughly minimize *E*(*z*). The random registration *F*^*t*^ = map(*Z*(*t*)) will belong to *MAP* and verify *Z*(*t*) = range(*F*^*t*^), so that *E*(*Z*(*t*)) = *E*(range(*F*^*t*^)) = cost(*F*^*t*^)). Consequently, for large *t*—with high probability—the random mapping *F*^*t*^ = map(*Z*(*t*)) will have a value of the cost functional cost(*F*^*t*^) close to minimize _*f* ∈*MAP*_ cost(*f*).

### 12.3. Cliques of Interactive Neurons

The BM energy function *E*(*z*) just defined turns out to involve only three sets of small cliques: **i)** *CL*_1_ is the set of all singletons *K* = {*i*}, with *i* = 1 … *N*. **ii)** *CL*_2_ is the set of all pairs *K* = {*i, j*} such that cells *b*_*i*_ and *b* _*j*_ are *neighbors* in *B*. **iii)** *CL*_3_ is the set of all triplets *K* = {*i, j, k*} such that cells *b* _*j*_ and *b*_*k*_ are both *neighbors* of *b*_*i*_ in *B*. Denote *CLQ* = *CL*_1_ ∪ *CL*_2_ ∪ *CL*_3_ the set of all cliques for our BM.

*Cliques in CL*_1_.. For each clique *K* = {*i*} in *CL*_1_, and each *z* ∈ *CONF*, define its energy Jmatch_*K*_(*z*) = Jmatch_*K*_(*z*_*i*_) by

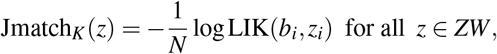

where LIK is given by (9). Set Jmatch_*K*_ ≡ 0 for *K* in *CL*_2_ ∪ *CL*_3_. For all *z* ∈ *CONF*, define the energy Ematch(*z*) by

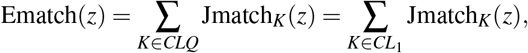

which implies that the registration *f* = map(*z*) verifies

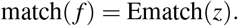

*Cliques in CL*_2_.. For all *z* ∈ *CONF*, all cliques *K* = {*i, j*} in *CL*_2_, define the clique energies Jover_*K*_(*z*) = Jover_*K*_(*z*_*i*_, *z* _*j*_) and Jstab_*K*_(*z*) = Jstab_*K*_(*z*_*i*_, *z* _*j*_) by 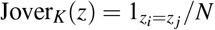 and

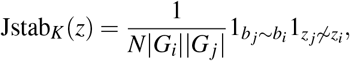

where |*G*_*i*_| and |*G*_*j*_| are the numbers of neighbors in *B* for cells *z*_*i*_ and *z* _*j*_, respectively. Set Jover_*K*_ = Jstab_*K*_ ≡ 0 for *K* in *CL*_1_ ∪ *CL*_3_. Define the two energy functions

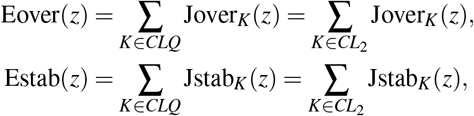

which implies that *f* = map(*z*) verifies over(*f*) = Eover(*z*) and stab(*f*) = Estab(*z*).

*Cliques in CL*_3_.. For each clique *K* = {*i, j, k*} in *CL*_3_, define the clique energy Jflip_*K*_ by

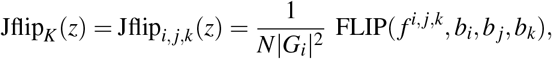

where *f* ^*i, j,k*^ is any registration mapping *b*_*i*_, *b* _*j*_, *b*_*k*_ onto *z*_*i*_, *z* _*j*_, *z*_*k*_. The indicator FLIP was defined in Sec. 11.2. Set Jflip_*K*_ ≡ 0 for *K* in *CL*_1_ ∪ *CL*_2_. Define the energy

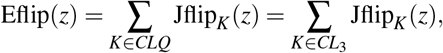

which implies that *f* = *F*(*z*) verifies flip(*f*) = Eflip(*z*).

Finally, define the clique energy *J*_*K*_ for all *K* ∈ *CLQ* by the linear combination

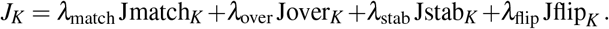

Summing this relation over all *K* ∈ *CLQ* yields

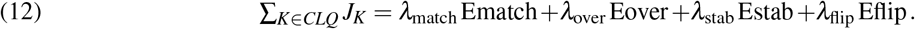

Define then the final BM energy function *z* ↦ *E*(*z*) by

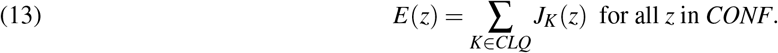

For any *z* ∈ *CONF*, the associated registration *f* = map(*z*) verifies match(*f*) = Ematch(*z*), over(*f*) = Eover(*z*), stab(*f*) = Estab(*z*), flip(*f*) = Eflip(*z*). By weighted linear combination of these equalities, and due to (12), we obtain for all configurations *z* ∈ *CONF, E*(*z*) = cost(*f*) when *f* = map(*z*) or, equivalently, when *z* = range(*f*).

### 12.4. Test Set of 100 Synthetic Image Pairs

As shown above, the minimization of cost(*f*) over all registrations *f* ∈ *MAP* is equivalent to seeking BM configurations *z* ∈ *CONF* with minimal energy *E*(*z*). We have implemented this minimization of *E*(*z*) by the long term asynchronous dynamics of the BM just defined. This algorithm was designed for the registration of image pairs exhibiting no cell division, and was, therefore, implemented after the automatic reduction of the generic registration problem, as indicated earlier. We have tested this specialized registration algorithm on a set BENCH_100_ of 100 pairs of successive images of simulated cell colonies exhibiting no cell divisions. These 100 image pairs were extracted from the benchmark set BENCH6 of synthetic image sequence described in section Sec. 2. The 100 pairs of cell sets *B, B*_+_ had sizes *N* = card(*B*) = card(*B*_+_) ranging from 80 to 100 cells.

For each test pair *B, B*_+_, each target window *W* (*j*) typically contained 30 to 40 cells. The set *CONF* of configurations had huge cardinality ranging from 10^130^ to 10^160^. But the average number of neighbors of a cell was around 4 to 5.

### 12.5. Implementation of BM minimization for cost(*f*)

The numbers *clq*_1_, *clq*_2_, *clq*_3_ of cliques in *CL*_1_, *CL*_2_, *CL*_3_ have the following rough ranges 80 ≤ *clq*1 ≤ 100, 160 ≤ *clq*_2_ ≤ 250, and 450 ≤ *clq*_3_ ≤ 600. For *k* = 1, 2, 3, denote *val*(*k*) the numbers of non-zero values for *J*_*K*_(*z*) when *z* runs through *CONF* and *K* runs through all cliques of cardinality *k*. One easily checks the rough upper bounds *val*(1) < 4.00e3; *val*(2) < 2.00e5; *val*(3) < 3.00e5. Hence, to automatically register *B* to *B*_+_, one could pre-compute and store all the possible values of *J*_*K*_(*z*) for all cliques *K* ∈ *CL*_1_ ∪ *CL*_2_ ∪ *CL*_3_ and all the configurations *z* ∈ *CONF*. This accelerates the key computing steps of the asynchronous BM dynamics, namely, for the evaluation of energy change Δ*E* = *E*(*z′*) − *E*(*z*), when configurations *z* and *z′* differs at only one site *j* ∈ *S*. Indeed, the single site modification 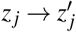 affects only the energy values *J*_*K*_(*z*) for the very small number *r*(*j*) of cliques *K*, which contain the site *j*. In our benchmark sets of synthetic images, one had *r*(*j*) < 24 for all *j* ∈ *S*. Hence, the computation of Δ*E* was fast since it requires retrieving at most 24 pairs of pre-computed *J*_*K*_(*z*), *J*_*K*_(*z′*), and evaluating the 24 differences *J*_*K*_(*z′*) − *J*_*K*_(*z*). Another practical acceleration step is to replace the ubiquitous computations of probabilities *p*(*t*) = exp(−*D/Temp*(*t*) by simply testing the value −*D/Temp*(*t*) against 100 precomputed logarithmic thresholds.

In our implementation of ABM dynamics, we used virtual temperature schemes such as *Temp*(*t*) = 50 · *ρ*^*t*^ with 0.995 ≤ *ρ* ≤ 0.999. The BM simulation was stopped when the stochastic energy *E*(*Z*(*t*)) had remained roughly stable during the last *N* steps. Since all target windows *W* (*j*) had cardinality smaller 40, the initial configuration *Z*(0) = *x* was computed via

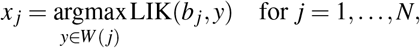

where the likelihoods LIK were defined by (9).

### 12.6. Weight Calibration

For the pair of successive *synthetic* images *J, J*_+_ displayed in Fig. 4, we have *N* = card(*B*) = card(*B*_+_) = 513 cells. The ground truth registration *f* is known by construction; we used it to apply the weight calibration described in Sec. 5. We set the meta-parameter *γ* to 1e10 and obtained the vector of weights

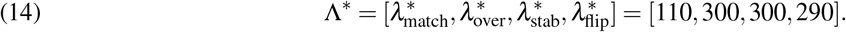

These weights are *kept fixed for all the 100 pairs* of images in the set BENCH_100_. The determined weights are used in the cost function cost(*f*) defined above. This correctly parametrized the BM energy function *E*(*z*). We then simulated the BM stochastic dynamics to minimize the BM energy *E*(*Z*(*t*)).

### 12.7. BM Simulations

We launched 100 simulations of the asynchronous BM dynamics, one for each pair of successive images in our test set BENCH_100_. For each such pair, the ground truth mapping *f* : *B* → *B*_+_ was known by construction and the stochastic minimization of the BM energy generated an estimated cells registration *f′* : *B* → *B*_+_. For each pair *B, B*_+_ in BENCH_100_, the accuracy of this automatically computed registration *f′* was evaluated by the percentage of cells *b* ∈ *B* such that *f′* (*b*) = *f* (*b*). When card(*B*) = *N*, our BM has *N* stochastic neurons, and the asynchronous BM dynamics proceeds by successive *epochs*. Each epoch is a sequence of *N* single site updates of the BM configuration. For each one of our 100 simulations of BM asynchronous dynamics, the number of epochs ranged from 250 to 450.

The average computing time was about eight seconds per epoch on a standard laptop, which entailed a computing time ranging from 30 to 50 minutes for each one of our 100 automatic registrations *f′* : *B* → *B*_+_. Note that the temperature scheme had not been optimized yet, so that these computing times are upper bounds. Earlier SBM studies [13, 15] indicate that the same energy minimizations on GPUs could provide a computational speedup by a factor ranging between 30 and 50.

We report registration accuracies in Tab. 2. For each pair of images in BENCH_100_, the accuracy of automatic registration was larger than 94.5%. The overall average registration accuracy was quite high at 99%.

**Table 2.**
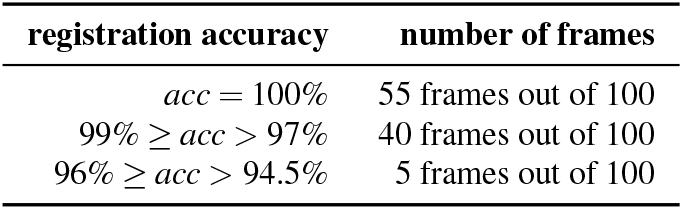
Registration accuracy for synthetic image sequence BENCH_100_. We consider 100 pairs of consecutive synthetic images (image sequence BENCH_100_). Automatic registration was implemented by BM minimization of the cost function cost(f), which was parametrized by the vector of optimized weights Λ∗ in (14). The average registration accuracy was 99%.

## 13. Registration for Cell Dynamics Involving Growth, Motion, and Cell Divisions

### 13.1. Tests of Cell Registration algorithms on Synthetic Data

We now consider more generic long synthetic image sequences of simulated cell colonies, with a small interframe duration of one minute. We still impose the mild constraint that no cell is lost between two successive images. The main difference with the earlier benchmark BENCH_100_ is that cells are *allowed to freely divide* during interframes, as well as to grow and to move. For the full implementation on 100 pairs of successive images, we first execute the parent-children pairing, and remove the identified parent-children triplets; we can then apply our cell registration algorithmic on the reduced sets cells. Our image sequence contained 760 true parent-children triplets, which we automatically identified with an accuracy of 100%. As outlined earlier, we removed all these identified cell triplets and then applied our tracking algorithm. This left us with a total of 1.26e4 cells (spread over 100 frames). Full automatic registration was then implemented with an accuracy higher than 99.5%.

### 13.2. Tests of Cell Registration algorithms on Laboratory Image Sequences

To test our cell tracking algorithm on pairs of consecutive images extracted from recorded image sequences of bacterial colonies, we had to automatically delineate all individual cells in each image. We use the Watershed algorithm [31] (also used, e.g., in [53]) to segment each frame into individual image segments containing one single cell each. In a second step, we then identified the contour of each single cell *b* by applying the Mumford-Shah algorithm [37, 68] within the image segment containing a cell *b*.

We then apply ad hoc nonlinear filters to remove minor segmentation artifacts. Since this procedure is quite time consuming for large images, we have implemented it to produce a training set of delineated individual cells to train a CNN for image segmentation. After automatic training, this CNN substantially reduces the runtime of the cell segmentation/delineation procedure. We show the resulting segmentations in Fig. 6.

**Fig. 6.**
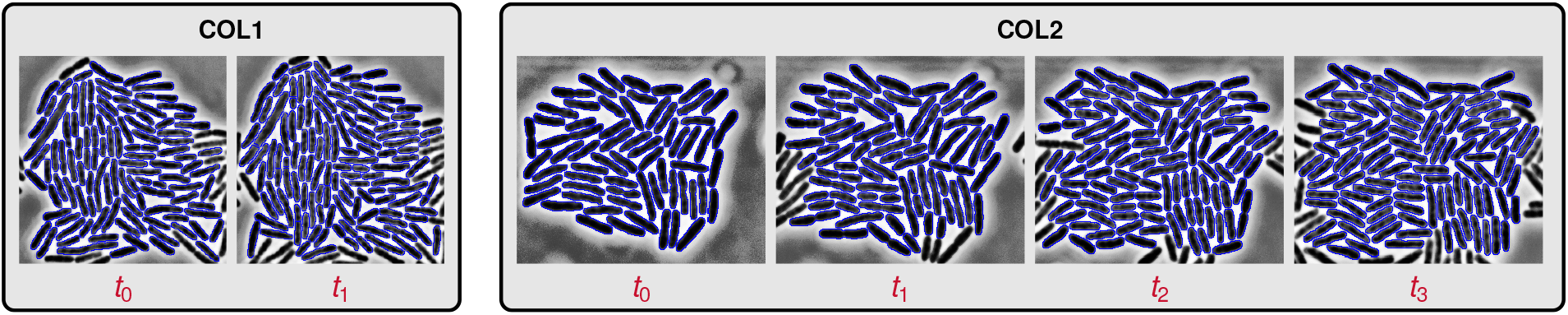
Segmentation results for experimental recordings of live cell colonies. We show two short image sequences extracts COL1 (left) and COL2 (right). The interframe duration is six minutes. The image sequence extract COL1 has only two successive image frames. The image sequence extract COL2 has four successive image frames. We are going to automatically compute four cell registrations, one for each pair of successive images in COL1 and COL2.

After each cell has been identified (i.e., segmented out) in each pair *J, J*_+_ of successive images, we transform *J, J*_+_ into binary images, where cells appear in white on a black background. For each resulting pair *B, B*_+_ of successive sets of cells, we apply the parent-children pairing algorithm outlined in Sec. 6 to identify all the short lineages. For the two successive images in COL1, the discovered short lineages are shown in Fig. 7 (left pair of images). Here, color designates the cell triplet algorithmically identified: parent cell in image *J* and its two children in image *J*_+_. We then remove each identified “*parent*” from *B* and its two children from *B*_+_. This yields the reduced cell sets *redB* and *redB*_+_. We can then apply our tracking algorithm (see 10) dedicated to situations where cells do not divide during the interframe.

**Fig. 7.**
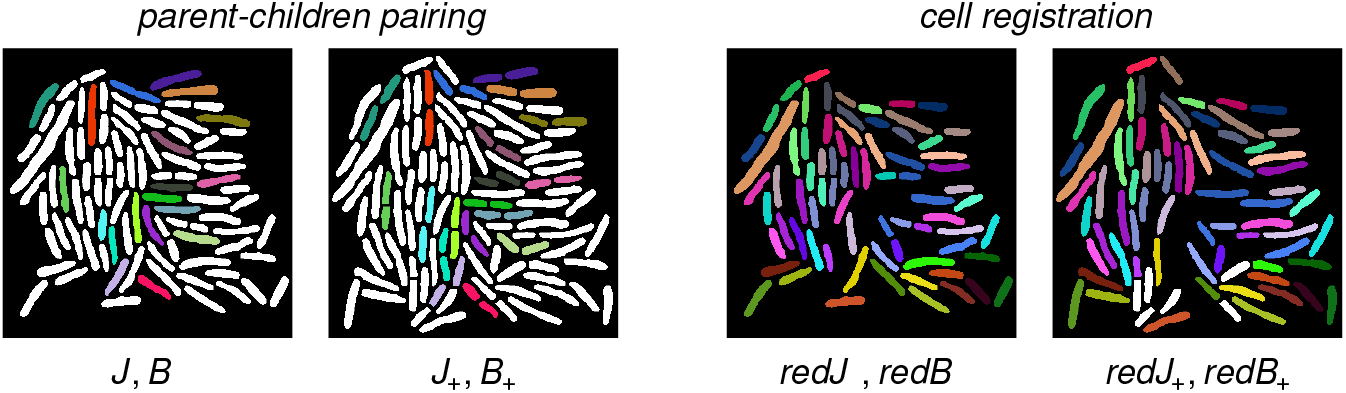
Cell tracking results for the pair COL1 of successive images J, J_+_ shown in Fig. 6. The interframe duration is six minutes. Left: Results for parent-children pairing on COL1. Automatically detected parent-children triplets are displayed in the same color. Right: Computed registration. The removal of the automatically detected parent-children triplets (see left column) generates the reduced cell sets redB and redB_+_. Automatic registration of redB and redB_+_ is again displayed via identical color for the registered cell pairs (b, b_+_). Mismatches are mostly due to previous errors in parent-children pairing (see Fig. 8 for a more detailed assessment).

For image sequences of live cell colonies we had to re-calibrate most of our weight parameters. The weight parameters used for these image sequences are summarized in Tab. 3.

**Table 3.** Cost function weights for parent-children pairing in the COL1 images displayed in Fig. 6.

| Weights | $\lambda_{\text{cen}}$ | $\lambda_{\text{siz}}$ | $\lambda_{\text{ang}}$ | $\lambda_{\text{gap}}$ | $\lambda_{\text{dev}}$ | $\lambda_{\text{rat}}$ | $\lambda_{\text{rank}}$ | $\lambda_{\text{over}}$ |
| --- | --- | --- | --- | --- | --- | --- | --- | --- |
| Value | 3 | 7 | 100 | .8 | 4 | .01 | .01 | 600 |

The BM temperature scheme was *Temp*(*t*) = 2000 (0.995)^*t*^, with the number of epochs capped at 5000. We illustrate our COL1 automatic registration results in Fig. 7 (right pair of images). Here, if cell *b* ∈ *redB* has been automatically registered onto cell *b*_+_ ∈ *redB*_+_, *b, b*_+_ share the same color. The cells colored in white in *redB*_+_ are cells which the registration algorithm did not succeed in matching to some cell in *redB*. These errors can essentially be attributed to errors in the parent-children pairing step. By visual inspection we have determined that there are 14 true parent-children triplets in the successive images of COL1. Our parent-children pairing algorithm did correctly identify 11 of these 14 triplets.To check further the performance of our registration algorithm on live images, we also report automatic registration results for “*manually prepared*” true versions of *redB* and *redB*_+_, obtained by removing “*manually*” the true parent-children triplets determined by visual inspection. For the short image sequence COL2, results are displayed in Fig. 8.

**Fig. 8.**
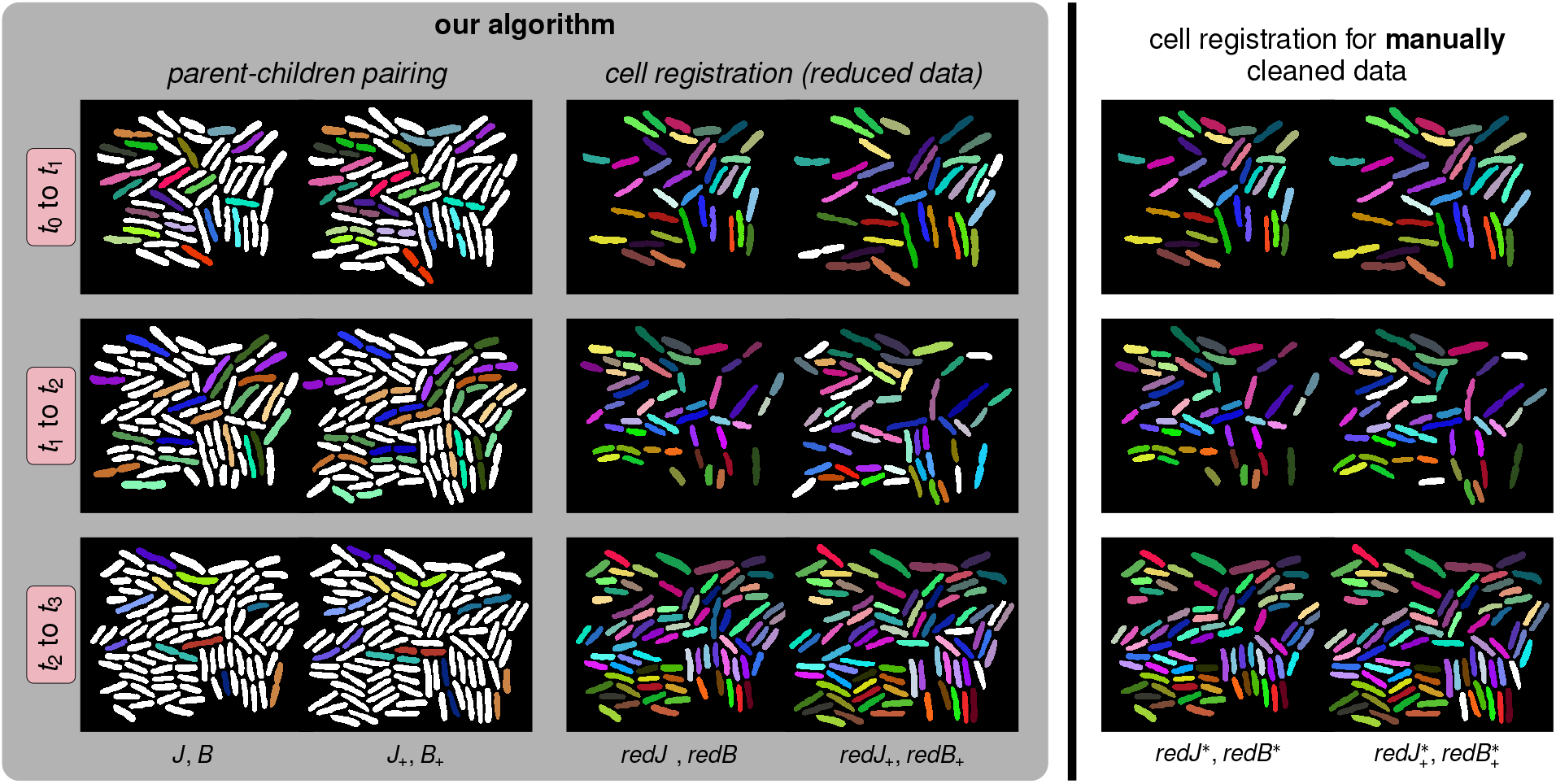
Cell tracking results for the short image sequence COL2 in Fig. 6. The interframe duration for COL2 is six minutes. COL2 involves four successive images J(t_i_), i = 0, 1, 2, 3. In our figure, each one of the three rows displays the automatic cell registration results between images J(t_i_) and J(t_i+1_) for i = 0, 1, 2. We report the accuracies of parent-children pairing and of the registration in Tab. 4. Left column: Results for parent-children pairing. Each parent-children triplet is identified by the same color for each parent cell an its two children. Middle column: Display of the automatically computed registration after removing the parent-children triplets already identified in order to generate two reduced sets redB and redB_+_ of cells. Again, the same color is used for each pair of automatically registered cells. The white cells in redB_+_ are cells which could not be registered to some cell in redB. Right column: To differentiate between errors induced during automatic identification of and errors generated by automatic registration between redB and redB_+_, we manually removed all “true” parent-children triplets and then applied our registration algorithm to this “cleaned” (reduced) cell sets redB^∗^ and 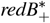.

**Table 4.**
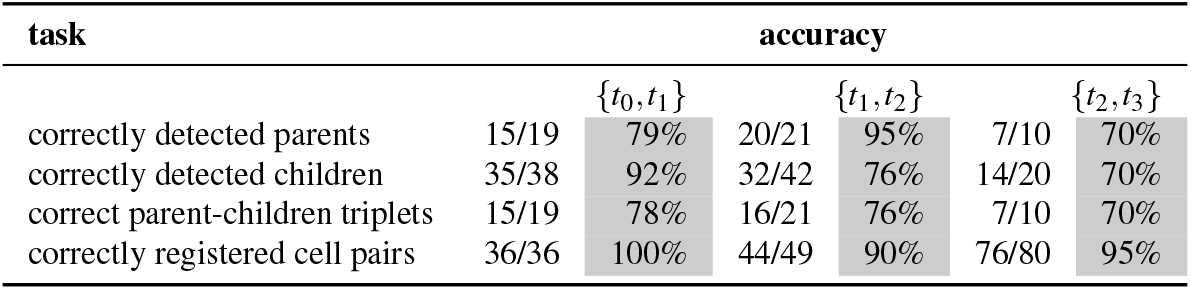
Cell tracking accuracy for the short image sequence COL2 in Fig. 6 with an interframe of six minutes. We report the ratio of correctly predicted cell matches over the total number of true cell matches and the associated percentages. The accuracy results quantify four distinct percentages of correct detections (i) for parent cells in image J, (ii) for children cells in image J_+_, (iii) for parent-children triplets, and (iv) for registered pairs of cells (b, b_+_) ∈ redB × redB_+_.

The display setup is the same: The left column shows the results of automatic parent-children pairing. The middle column illustrates the computed registration after automatic removal of the computer identified parent-children triplets. The third column displays the computed registration after removing “*manually*” the true parent-children triplets determined by visual inspection. Note that the overall matching accuracy can be improved if we reduce errors in the parent-children pairing. We report quantitative accuracies in Tab. 4. For parent-children pairing, accuracy ranges between 70% and 78%. For pure registration after correct parent-children pairing, accuracy ranges between 90% and 100%.

## 14. Conclusions and Future Work

We have developed a methodology for automatic cell tracking in recordings of dense bacterial colonies growing in a mono-layer. We have also validated our approach using synthetic data from agent based simulations, as well as experimental recordings of *E. coli* colonies growing in microfluidic traps. Our next goal is to streamline our implementation for systematic cell registration on experimentally acquired recordings of such cell colonies, to enable automated quantitative analysis and modeling of cell population dynamics and lineages.

There are a number of challenges for our cell tracking algorithm: Inherent imaging artifacts such as noise or intensity drifts, cells overlaps, similarity of cell shape characteristics across the population, tight packing of cells, somewhat large interframe times, cell growth combined with cell motion and cell divisions, represent just a few of these challenges. Overall, the cell tracking problem has combinatorial complexity, and for large frames is beyond the concrete patience of human experts. We tackle these challenges by developing a two-stage algorithm that first identifies parent-children triplets and subsequently computes cell registration from one frame to the next, after reducing the two original cell sets by automatic removal of the identified parent-children triplets. Our algorithms specify innovative cost functions dedicated to these registration challenges. These cost functions have combinatorial complexity. To discover good registrations we minimize these cost functions numerically by intensive stochastic simulations of specifically structured BMs. We have validated the potential of our approach by reporting promising results obtained on long synthetic image sequences of simulated cell colonies (which naturally provide a ground truth for cell registration from one frame to the next). We have also successfully tested our algorithms on experimental recordings of live bacterial colonies.

In future work we will further improve the stability and accuracy of our cell registration algorithms by exploring natural modifications of our cost functions, in particular to improve the accuracies of our automatic detection of cell divisions, and to handle also the unavoidable new cell arrivals into or departures from the current field of vision. Moreover, we will work on our BM simulation algorithms to deploy their stochastic dynamics in semi synchronous formats on parallel computing platforms (in particular, graphic processing units) to drastically reduce their runtime.

## Acknowledgements

This work was partly supported by the National Science Foundation (NSF) through the grants DMS-1854853 (AM & RA), DMS-2009923 (AM & RA), 1662305 (KJ), MCB-1936770 (KJ), and DMS- 2012825 (AM); the joint NSF-National Institutes of General Medical Sciences Mathematical Biology Program grant DMS-1662290 (MRB); and the Welch Foundation grant C-1729 (MRB). Any opinions, findings, and conclusions or recommendations expressed herein are those of the authors and do not necessarily reflect the views of the NSF or the Welch Foundation. This work was completed in part with resources provided by the Research Computing Data Core at the University of Houston.

## Appendix A. Stochastic Dynamics of BMs

Notations and terminology refer to Sec. 7. Consider a BM network of *N* stochastic neurons *U*_*j*_, with finite configuration set *CONF* = *W* (1) × … × *W* (*N*). At time *t*, let *Z*_*j*_(*t*) ∈ *W* (*j*) be the random state of neuron *U*_*j*_, and the BM configuration *Z*(*t*) ∈ *CONF* is then *Z*(*t*) = {*Z*_1_(*t*), …, *Z*_*N*_(*t*)}. Fix as in Sec. 7 a sequence *Temp*(*t*) of virtual temperatures slowly decreasing to 0 for large *t*.

There are two main options to implement the Markov chain dynamics *Z*(*t*) → *Z*(*t* + 1) (see [10]).

### Asynchronous BM Dynamics

Generate a long random sequence of sites *m*(*t*) ∈ *S* = {1, …, *N*}, for instance by concatenating successive random permutations of the set *S*. At time *t*, the only neuron which may modify its current state is *U*_*m*(*t*)_. For brevity, write *M* = *m*(*t*). The neuron *U*_*M*_ will compute its new random state *Z*_*M*_(*t* + 1) ∈ *W* (*M*) by the following *updating procedure*:

- For each *y* in *W* (*M*), define a new configuration *Y* ∈ *CONF* by *Y*_*M*_(*t*) = *y*, and *Y*_*j*_(*t*) = *Z*_*j*_(*t*) for all *j* ≠ *M*. Let Δ(*y*) = *E*(*Y*) − *E*(*Z*(*t*)) be the corresponding BM energy change.
- In the finite set *W* (*M*), select any *z* such that Δ(*z*) = min_*y*∈*W*(*M*)_ Δ(*y*), and set *D* = max {0, Δ(*z*)}
- Compute the probability *p* = exp(−*D/Temp*(*t*)).
- The new random state *Z*_*M*_(*t* + 1) of neuron *U*_*M*_ will be equal to *z* with probability *p* and equal to the current state *Z*_*M*_(*t*) with probability 1 − *p*.
- For all *j* ≠ *M*, the new state *Z*_*j*_(*t* + 1) of neuron *U*_*j*_ remains equal to its current state *U*_*j*_(*t*)

### Synchronous BM Dynamics

Fix a *synchrony* parameter 0 < *α* < 1, usually around 50%. At each time *t, all* neurons *U*_*j*_ synchronously, but independently compute their own random *binary tag tag*_*j*_ (*t*), equal to 1 with probability *α*, and to 0 with probability (1 − *α*). Let *SYN*(*t*) be the set of all neurons. All the neurons *U*_*j*_ such that *tag*_*j*_ (*t*) = 1 then synchronously and independently compute their new random states *Z*_*j*_(*t* + 1) ∈ *W* (*j*) by applying the updating procedure given above. And for all *j* such that *tag*_*j*_ (*t*) = 0, the new state *Z*_*j*_(*t* + 1) of *U j* remains equal to *Z*_*j*_(*t*).

### Comparing Asynchronous and Synchronous BM Dynamics

As *t* becomes large, and for temperatures *Temp*(*t*) slowly decreasing to 0, both BM dynamics generate with high probability configurations *Z*(*t*) which provide deep local minima *E*(*Z*(*t*)) of the BM energy function. The asynchronous dynamics can be fairly slow. But the synchronous dynamics is much faster since it emulates efficient forms of *parallelel simulated annealing* (see [11, 75]) and is directly implementable on GPUs.

## Footnotes

* Cell Tracking Challenge: http://celltrackingchallenge.net (accessed 03/2021).

